# *In Silico* Analysis Predicting Effects of Deleterious SNPs of Human *RASSF5* Gene on its Structure and Functions

**DOI:** 10.1101/2020.06.05.952952

**Authors:** Md. Shahadat Hossain, Arpita Singha Roy, Md. Sajedul Islam

## Abstract

Ras association domain-containing protein 5 (RASSF5), one of the prospective biomarkers for tumors, generally plays a crucial role as a tumor suppressor. As deleterious effects can result from functional differences through SNPs, we sought to analyze the most deleterious SNPs of *RASSF5* as well as predict the structural changes associated with the mutants that hamper the normal protein-protein interactions. We adopted both sequence and structure based approaches to analyze the SNPs of RASSF5 protein. We also analyzed the putative post translational modification sites as well as the altered protein-protein interactions that encompass various cascades of signals. Out of all the SNPs obtained from the NCBI database, only 25 were considered as highly deleterious by six *in silico* SNP prediction tools. Among them, upon analyzing the effect of these nsSNPs on the stability of the protein, we found 17 SNPs that decrease the stability. Significant deviation in the energy minimization score was observed in P350R, F321L, and R277W. Besides this, docking analysis confirmed that P350R, A319V, F321L, and R277W reduce the binding affinity of the protein with H-Ras, where P350R shows the most remarkable deviation. Protein-protein interaction analysis revealed that RASSF5 acts as a hub connecting two clusters consisting of 18 proteins and alteration in the RASSF5 may lead to disassociation of several signal cascades. Thus, based on these analyses, our study suggests that the reported functional SNPs may serve as potential targets for different proteomic studies, diagnosis and therapeutic interventions.

## Introduction

Ras association domain-containing protein 5 (RASSF5) is the leading member of Ras effector super family protein that prevents tumor growth by facilitating G_1_/S arrest of cell cycle^1-6^. RASSF5 is a proapoptotic component of Ras and induces p53 mediated apoptosis^4^. RASSF5 is implicated in a range of cell reactions including apoptosis, senescence, cell cycle regulation, differentiation and cell proliferation^5^. Inactivation of RASSF5 is found to be associated in the oncogenesis, proliferation and weak diagnosis of human cancers^5^. In various tumor tissues, low *RASSF5* expression has been discovered and can therefore be considered as a prospective biomarker for tumors. RASSF5, in a range of human cancers, is shown to play crucial functions and is identified as a putative tumor suppressor^7^.

The complete length of *RASSF5* gene is 81 kb and it is located at the locus 1q32.1 that covers several isoforms resulting from alternative mRNA splicing and differential promoter usage, which may have different roles in oncogenesis^3^. NORE1A, with a molecular weight of around 47 kDa, is a bigger isoform of the *RASSF5A* gene that encodes a protein of 418 amino acids^8^. The RASSF5A protein structure includes an N-terminal proline-rich section containing prospective SH3-binding sites, and a nuclear localization signal, a cysteine-rich C1-type zinc finger (C1) domain, a Ras-association (RA) domain that interacts with GTP-bound H-Ras or several other Ras subfamily GTPases, and a Salvador-RASSF-Hippo^9^ domain that contains C-type carboxy-terminal tail (**Figure 1**), which is essential for binding to pro-apoptotic kinases MST1/215 and interaction factors WW45/SAV1^10,11^. While RASSF5 has a large degree of homology with other RASSF members, only the RASSF5A contains a single proline-rich motif (PPxY)^4^.

**Figure 1:**
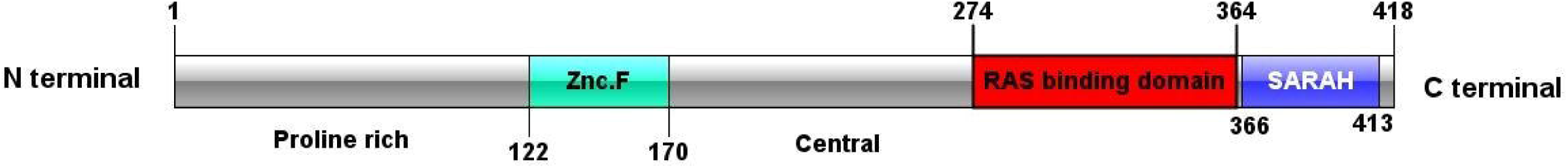
Protein structure of RASSF5 (Adopted and modified from *Shuofeng et. al*. 2018^83^)

Human genome shares about 99.9% identical DNA sequences globally and the rest 0.1% of the genome contains individual variations. This variation in the individual genome is resulted from random mutations^12^. Among them, the most common type of mutation is single nucleotide polymorphism (SNP), which is a single base substitution in the alleles^13^. SNPs occur throughout the genome at the rate of 1 in 1000 base pairs and the most significant SNPs are found in the coding regions^12^. Till now, about 500,000 SNPs are enlisted in the coding regions^13^. Amongst them, there are mainly two categories of SNPs observed, which are: Synonymous and Non-synonymous (nsSNP) SNPs. Synonymous SNPs refer to the conditions where mutations in the DNA sequences do not alter the amino acid sequences, whereas, non-synonymous SNPs, especially Missense SNPs, are responsible for amino acid substitutions and protein variations in human^14^. Previous studies show that nsSNPs are responsible for around 50 percent of the mutations involved in various genetic diseases^15,16^ along with several autoimmune and inflammatory diseases^17-19^.

Deleterious or neutral impacts on protein structure or function can result from functional differences through SNPs^20^. Detrimental effects consist of damage to protein structures and gene regulation^21^. Moreover, alterations in the protein sequence may ultimately affect alteration of protein charge, geometry, hydrophobicity^22^, dynamics, translation, and inter/intra protein interaction^6,23,24^ putting cells in danger^25^. This information confirms that nsSNPs, especially missense SNPs, are associated with various human diseases. Recent studies on the nsSNPs using computational approaches reveal the potential impact of mutation in understanding the molecular mechanisms of various diseases^26-28^. Structural and functional analysis of nsSNPs may facilitate the development of personalized medicine based on genomic variation. The impact of nsSNPs on RASSF5 protein regarding disease pathogenesis is not yet well-understood. Thus, in the present study, we explored various bioinformatics tools and servers to find out the functional effects of nsSNPs of RASSF5 protein.

## Results

### SNP annotation

We retrieved RASSF5 SNPs using NCBI dbSNP database that contained 37426 SNPs in intronic region, 215 SNPs in 5’UTR region, 1720 SNPs in 3’UTR region, and 757 SNPs in coding sequence. Out of the SNPs in coding sequence, 495 SNPs were missense (nsSNPs) and 236 SNPs were synonymous SNPs. For our current study, we only selected nsSNPs for further analyses as the change in codon results in different amino acids. Overview of the whole methodological approaches is summarized in a schematic diagram (**Figure 2**).

**Figure 2:**
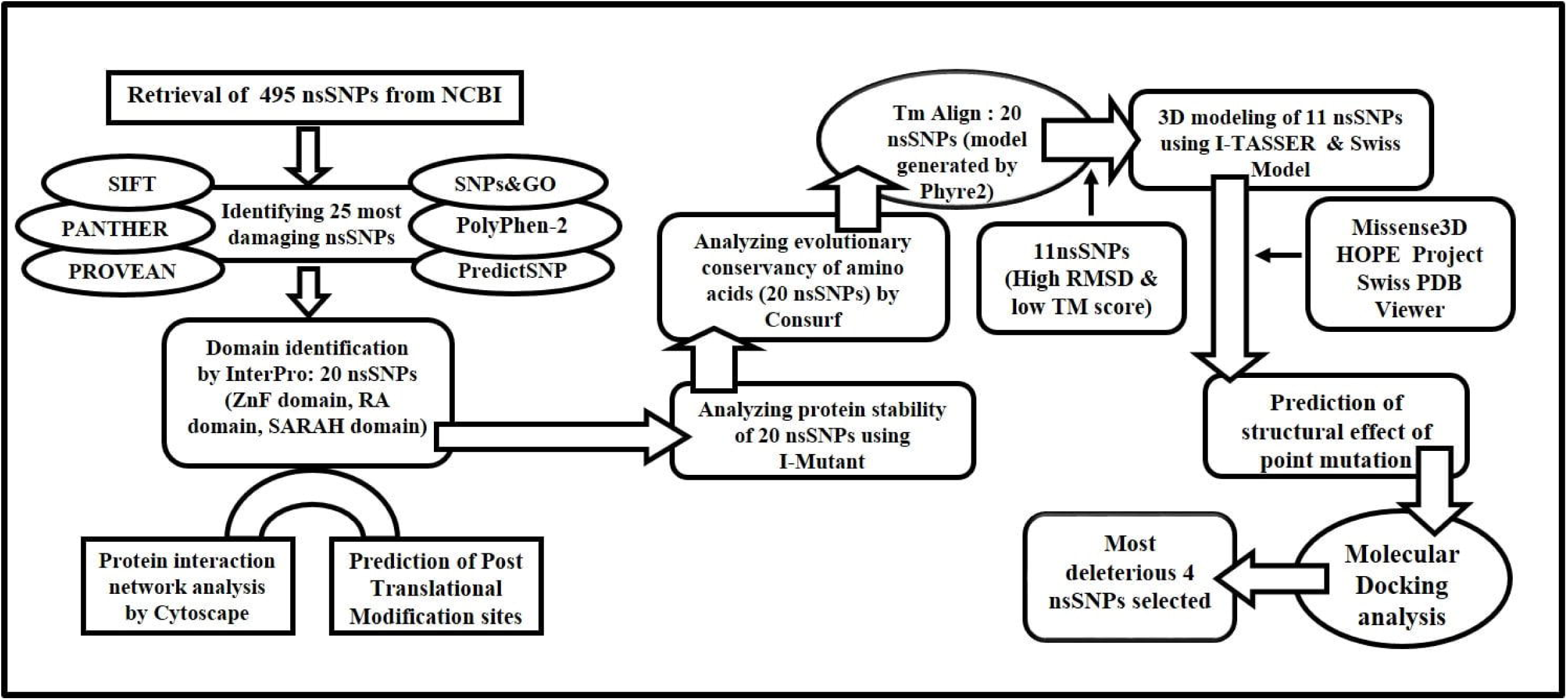
Schematic diagram summarizing the study

### Identification of deleterious nsSNPs

We used six different *in silico* nsSNP prediction tools (SIFT, PANTHER, PolyPhen-2, SNPs&GO, PROVEAN, and PredictSNP) to find out the deleterious SNPs that can significantly alter the structure or function of RASSF5 protein. Out of 495 nsSNPs, a total of 25 were predicted to be deleterious SNPs in all computational algorithms (**Table 1**).

**Table 1:**
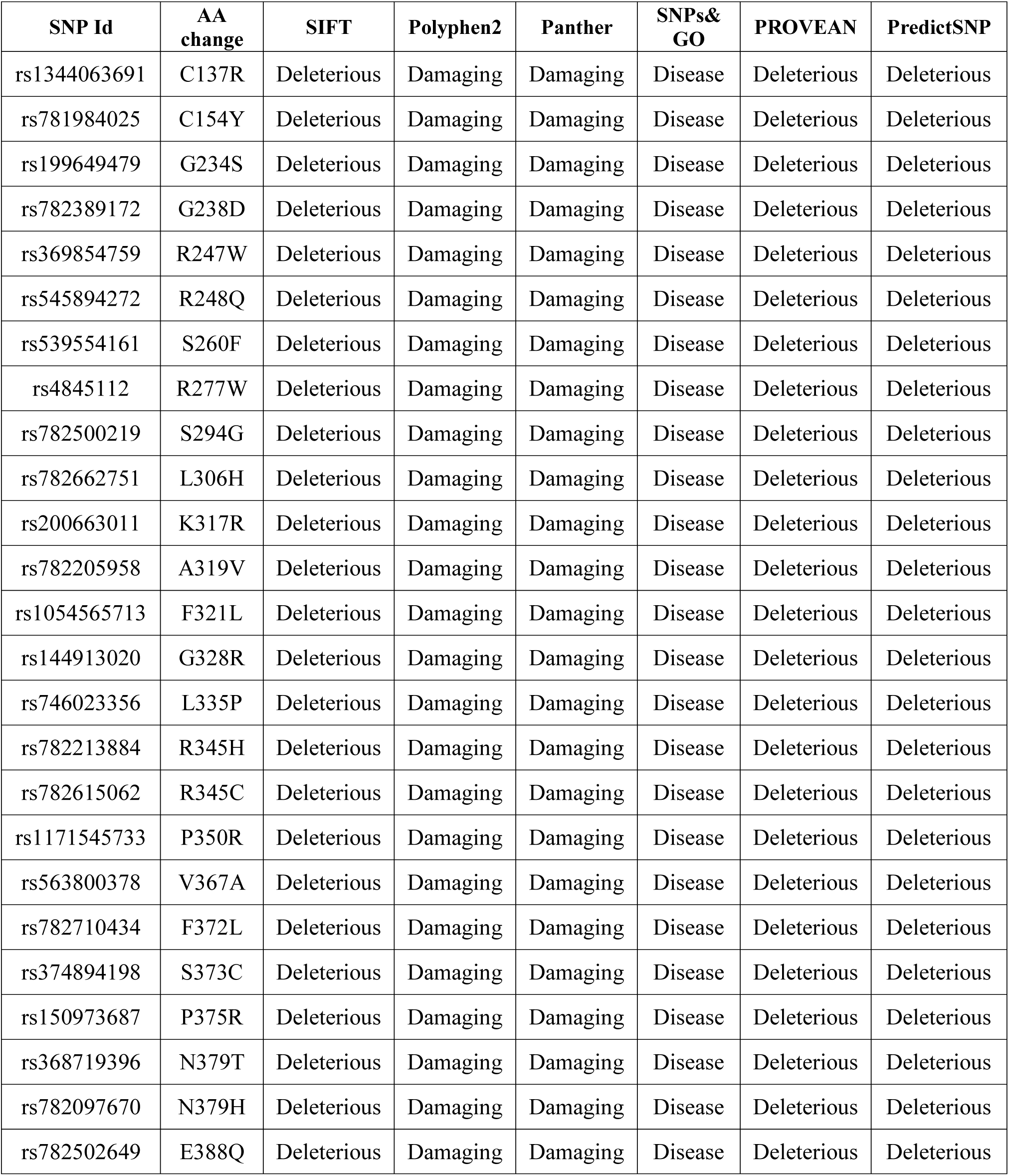
High risk nsSNPs identified by six *in silico* programs

| SNP Id | AA change | SIFT | Polyphen2 | Panther | SNPs&GO | PROVEAN | PredictSNP |
| --- | --- | --- | --- | --- | --- | --- | --- |
| rs1344063691 | C137R | Deleterious | Damaging | Damaging | Disease | Deleterious | Deleterious |
| rs781984025 | C154Y | Deleterious | Damaging | Damaging | Disease | Deleterious | Deleterious |
| rs199649479 | G234S | Deleterious | Damaging | Damaging | Disease | Deleterious | Deleterious |
| rs782389172 | G238D | Deleterious | Damaging | Damaging | Disease | Deleterious | Deleterious |
| rs369854759 | R247W | Deleterious | Damaging | Damaging | Disease | Deleterious | Deleterious |
| rs545894272 | R248Q | Deleterious | Damaging | Damaging | Disease | Deleterious | Deleterious |
| rs539554161 | S260F | Deleterious | Damaging | Damaging | Disease | Deleterious | Deleterious |
| rs4845112 | R277W | Deleterious | Damaging | Damaging | Disease | Deleterious | Deleterious |
| rs782500219 | S294G | Deleterious | Damaging | Damaging | Disease | Deleterious | Deleterious |
| rs782662751 | L306H | Deleterious | Damaging | Damaging | Disease | Deleterious | Deleterious |
| rs200663011 | K317R | Deleterious | Damaging | Damaging | Disease | Deleterious | Deleterious |
| rs782205958 | A319V | Deleterious | Damaging | Damaging | Disease | Deleterious | Deleterious |
| rs1054565713 | F321L | Deleterious | Damaging | Damaging | Disease | Deleterious | Deleterious |
| rs144913020 | G328R | Deleterious | Damaging | Damaging | Disease | Deleterious | Deleterious |
| rs746023356 | L335P | Deleterious | Damaging | Damaging | Disease | Deleterious | Deleterious |
| rs782213884 | R345H | Deleterious | Damaging | Damaging | Disease | Deleterious | Deleterious |
| rs782615062 | R345C | Deleterious | Damaging | Damaging | Disease | Deleterious | Deleterious |
| rs1171545733 | P350R | Deleterious | Damaging | Damaging | Disease | Deleterious | Deleterious |
| rs563800378 | V367A | Deleterious | Damaging | Damaging | Disease | Deleterious | Deleterious |
| rs782710434 | F372L | Deleterious | Damaging | Damaging | Disease | Deleterious | Deleterious |
| rs374894198 | S373C | Deleterious | Damaging | Damaging | Disease | Deleterious | Deleterious |
| rs150973687 | P375R | Deleterious | Damaging | Damaging | Disease | Deleterious | Deleterious |
| rs368719396 | N379T | Deleterious | Damaging | Damaging | Disease | Deleterious | Deleterious |
| rs782097670 | N379H | Deleterious | Damaging | Damaging | Disease | Deleterious | Deleterious |
| rs782502649 | E388Q | Deleterious | Damaging | Damaging | Disease | Deleterious | Deleterious |

### Identification of nsSNPs on the domains of RASSF5

InterPro, a domain identification tool, predicts the domains and active sites of a protein through the functional analysis of protein families. It predicted three functional domains of RASSF5, which are Zinc finger domain (122-170), Ras-association domain (274-364), and SARAH domain (366-413) and demonstrated that 20 out of 25 nsSNPs are positioned on these domains (**Figure S1**).

### Determination of protein structural stability

To predict the changes in the stability of RASSF5 in terms of RI and free energy change values (DDG), we used I-Mutant tool that introduced point mutation in the RASSF5 protein. The outcome revealed that 17 out of 20 deleterious nsSNPs decreased stability (**Table 2**).

**Table 2:**
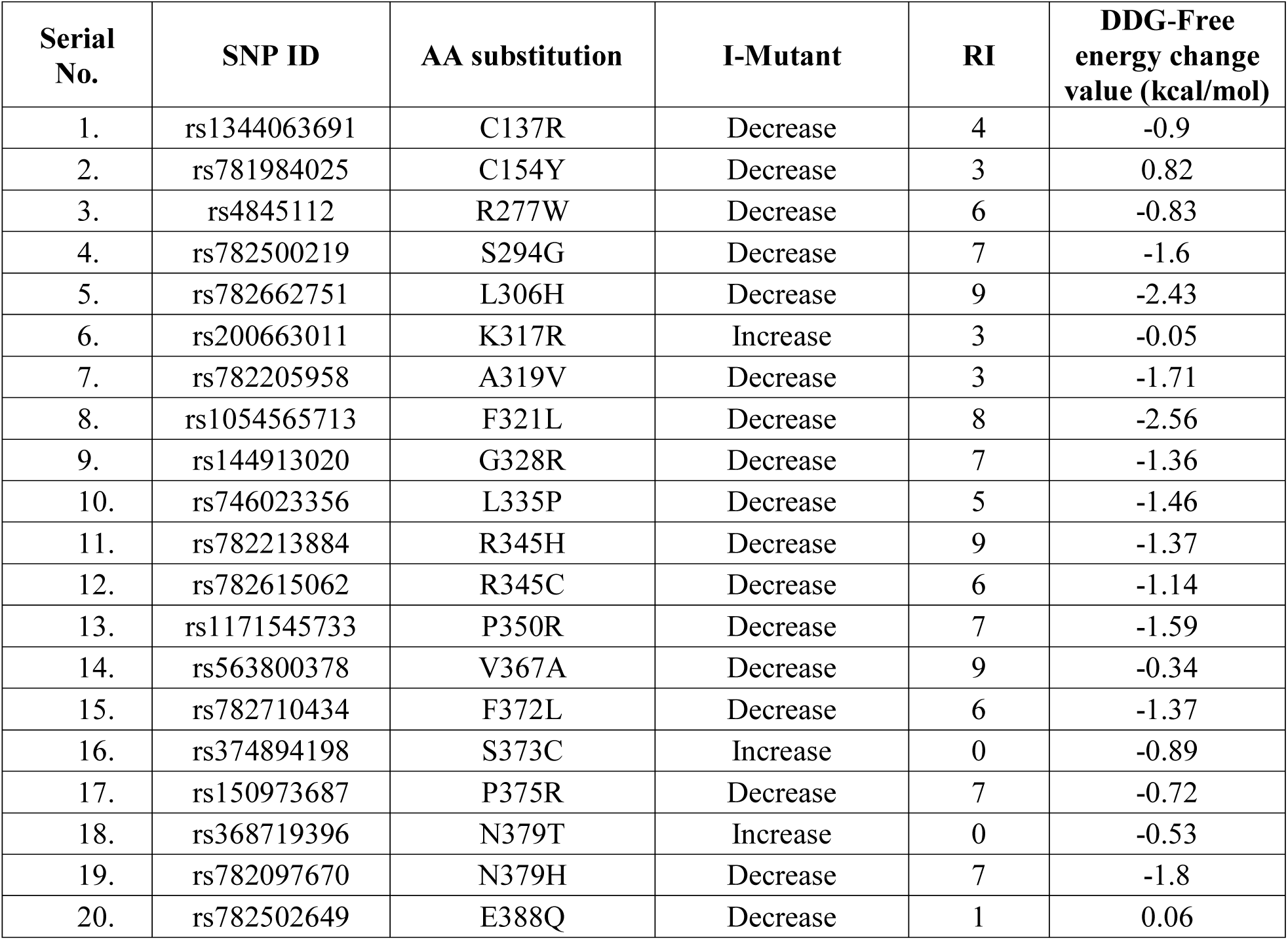
Effect of nsSNPs on protein stability predicted by I-MUTANT 2.0

### Evolutionary conservation analysis

The evolutionary conservancy of amino acid residues of the native RASSF5 was determined by ConSurf web server. It identified structural and functional residues of the 20 high risk nsSNPs of RASSF5 protein using evolutionary conservation and solvent accessibility. We found that R277, K317, P350, P375, N379, and E388 are exposed and functional whereas, C154, S294, L306, A319, L335, R345, V367, F372, and S373 residues are buried and structural. All 15 of these residues are highly conserved (**Table S1**). Additionally, G328 is predicted to be moderately conserved and exposed whereas, C137, and F321 are conserved and buried (**Figure 3**).

**Figure 3:**
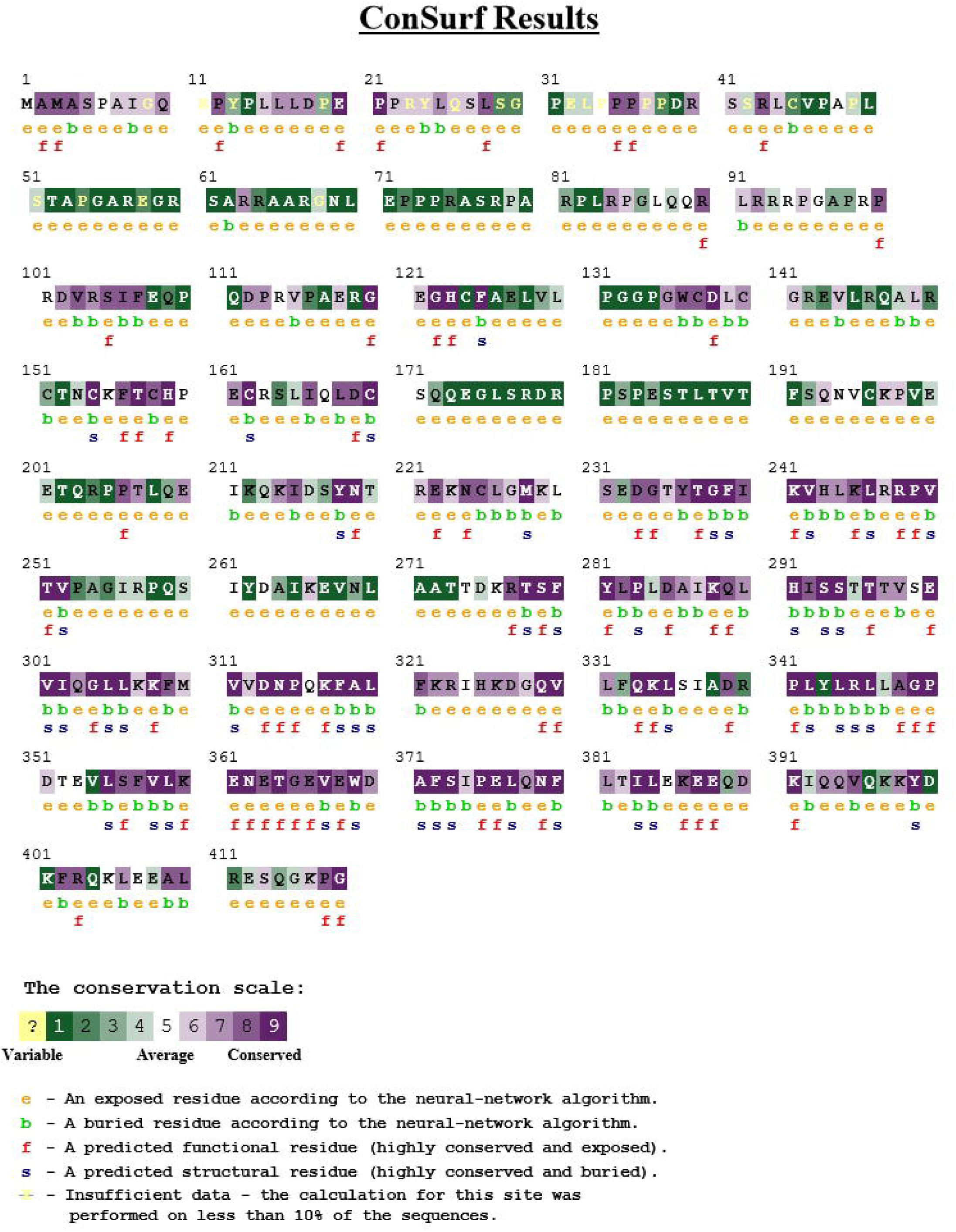
Evolutionary conservancy of RASSF5 produced by Consurf

### Structure analysis of wild type and mutant models

We used Phyre2 homology modeling tool to produce the 3D structure of native and 20 mutants of RASSF5 protein. We selected 1rfh (113-171), 3ddc (205-362), 4lgd (366-405) templates for zinc finger domain, Ras-binding domain and SARAH domain respectively using Phyre2. The 20 deleterious nsSNPs were individually replaced in the native sequence of the three templates and 3D models for all of the mutants were predicted.

Structural similarities between native and mutant models were investigated based on TM-score and RMSD scores using TM-align tool. The mutant models for the two nsSNPs found in zinc finger domain and seven nsSNPs found in SARAH domain showed high TM-score and low RMSD value in comparison with low TM-score and high RMSD value of the eleven nsSNPs found in Ras-binding domain (**Table S2**). As higher RMSD value indicates greater structural dissimilarity between wild type and mutant models, we considered the 11 nsSNPs found in the Ras-binding domain for further analyses. To substantiate the validity of this finding, we used I-TASSER for further structural analysis of these 11 nsSNPs in RASSF5 protein. Mutant models with lowest C-score were selected for superimposition over native structure.

Furthermore, 3D structures of the aforementioned 11 nsSNPs of RASSF5 protein were analyzed by SWISS MODEL to study protein solvation and torsion. The template used for the investigation of these nsSNPs was 3ddc.1.B. The result showed that values for solvation and torsion for wild type structure were -0.16Å^2^ and -1.03^0^ (degree) respectively but the values deviated significantly for the mutant structures (**Table S3**).

### Structural effect of point mutation on human RASSF5 protein

Project HOPE server revealed that the mutant residues of R277W and P350R with bigger sizes are more hydrophobic than the wild type residues and these variation in size and hydrophobicity can disrupt the H-bond interactions with the adjacent molecules. Moreover, P350 provides special backbone conformation but Arginine replacement may interrupt that formation (**Figure 4**). In case of R345H and R345C mutants, disruption of both salt bridge and hydrogen bond were observed. Flexibility and rigidity of a protein structure is essential for exhibiting specific function. Here, the analysis showed that amino acid substitution in G328R can disrupt the flexibility whereas introducing Glycine in S294G can disturb the required rigidity of RASSF5 protein. Besides this, L306P, K317R, A319V, F321L, L335P mutants exposed different properties and hence may significantly affect the functional Ras associating domain (**Table S4**).

**Figure 4:**
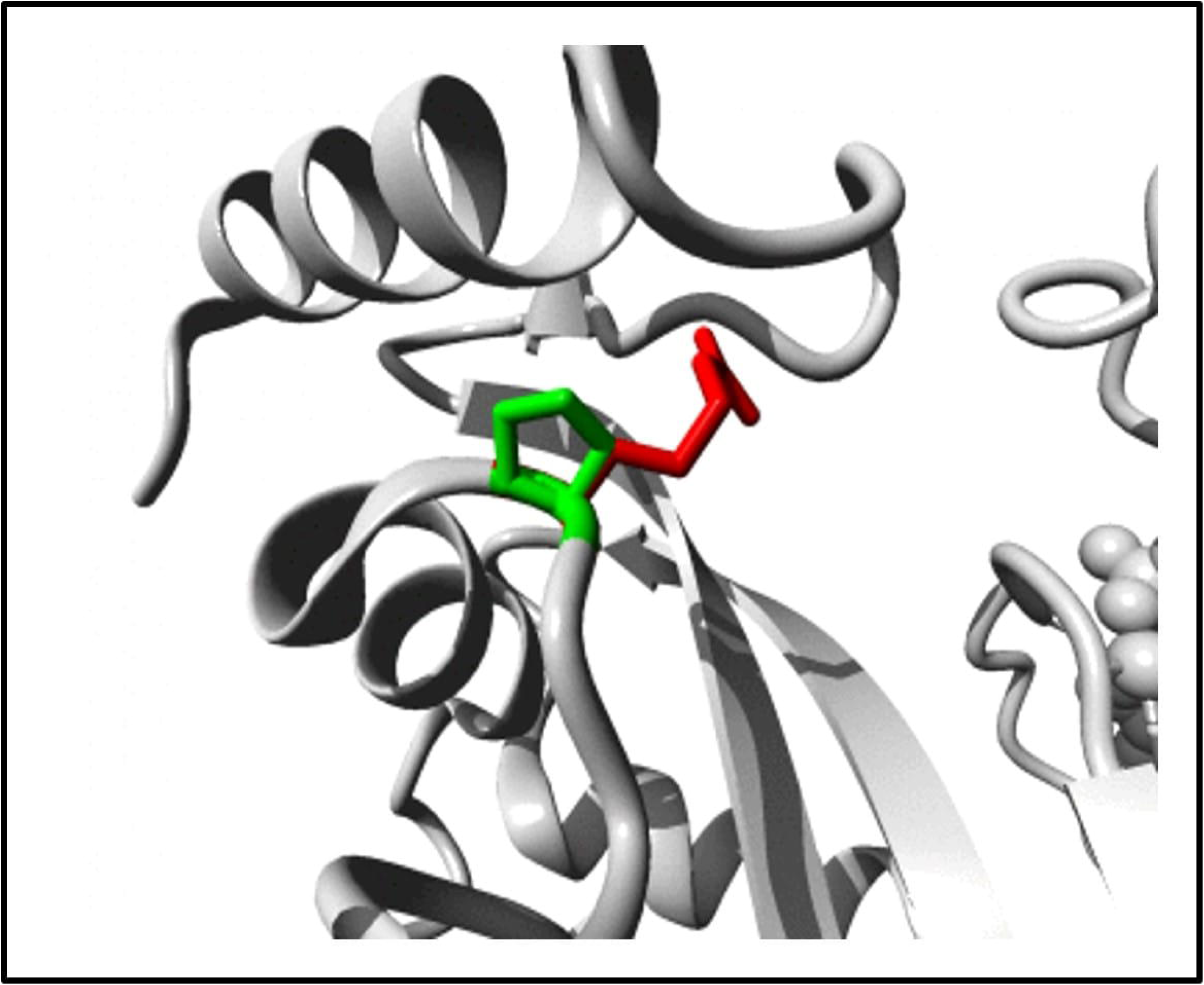
Structural alteration of the wild type residue P350 by the mutant R350 illustrated by Project Hope. The wild type residue is presented as green and the mutant residue is shown in red.

Missense3D tool presented that L335P variant introduced a buried Proline that provides constrained backbone conformation. Furthermore, R345H disrupted the side-chain and main-chain H-bond(s) formed by wild type buried Arginine residue and R345C substitution causes the enlargement of cavity volume by 98.064 L^3^ exceeding the cutoff value (**Table S5)**. Structural changes along with energy state was also observed in the 11 mutant models of RASSF5 protein obtained from Swiss PDB viewer mutation tool. The total energy of the wild type structure was found to be -17442.365 kJ/mol after energy minimization. Molecular simulation result showed that K317R (−17579.576kJ/mol) significantly decreased total energy whereas, P350R (−16379.468 kJ/mol) (**Figure 5**), F321L (−16381.121 kJ/mol), and R277W (−16448.330 kJ/mol) were the top three substitutions which increased total energy after energy minimization (**Table S6**).

**Figure 5:**
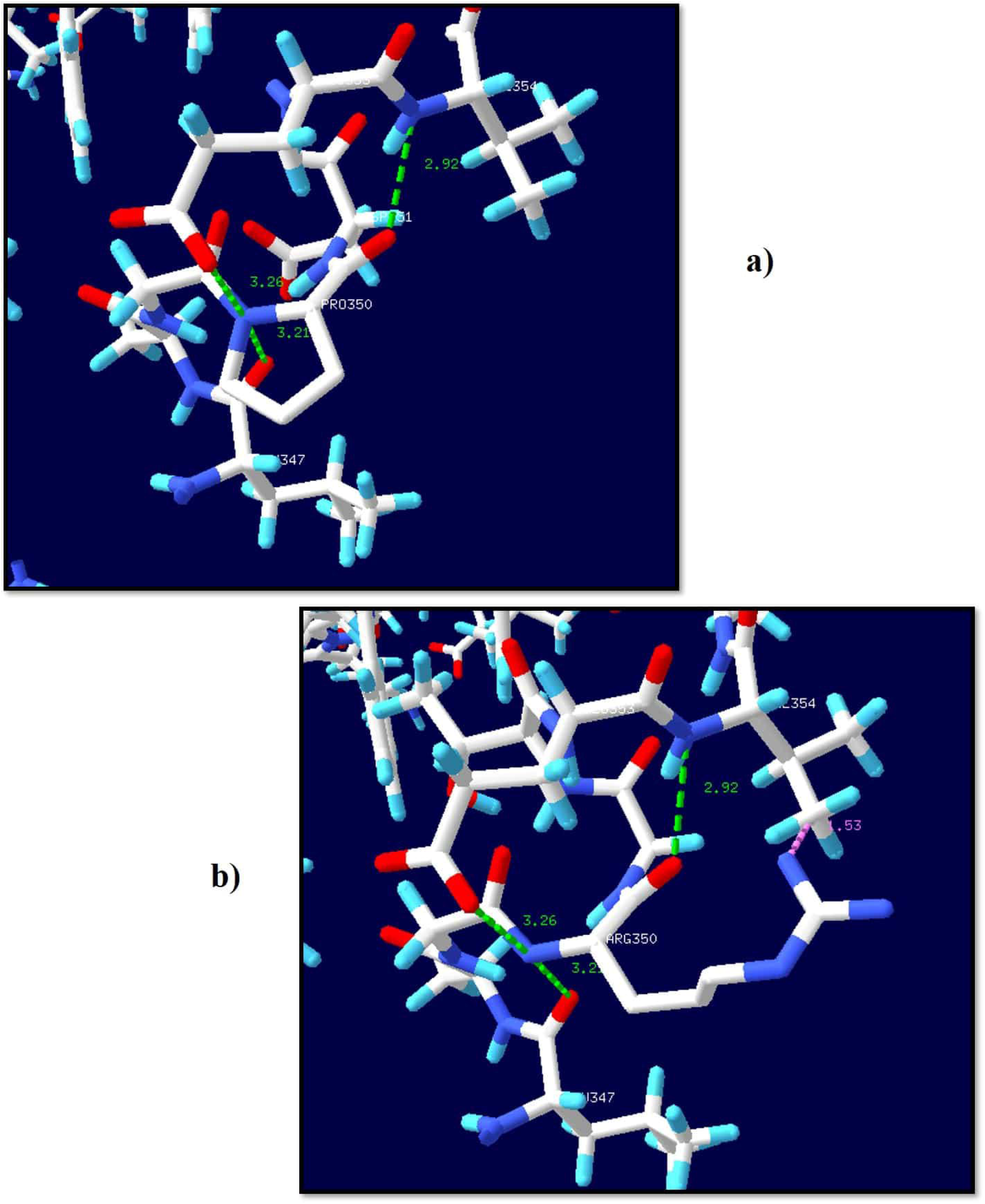
Structural Effect of P350R on the native structure of RASSF5 presented by Swiss PDB Viewer. **a)** P350 forms three H-bonds with T347 (3.21 Å), E353 (3.26 Å), and V354 (2.92 Å), which are indicated by green discontinuous lines. **b)** R350 forms a clash with V354 (1.53 Å) presented by pink discontinuous line along with the three H-bonds.

### Molecular docking analysis

Molecular docking studies represented the binding affinity of the mutated RASSF5 protein with H-Ras protein (**Figure 6**). Out of 11 nsSNPs, four (4) mutants (P350R, A319V, F321L, and R277W) significantly reduce the binding affinity with H-Ras protein (**Table 3**). The binding affinity (kcal/mol) and bonding interaction patterns of these docked complexes were studied using UCSF Chimera software. P350R variant disclosed the most significant reduction in binding affinity with H-Ras protein. The wild type peptide sequence forms five (5) hydrogen bonds with binding affinity -8.3 kcal/mol whereas the mutant forms three hydrogen bonds with the target protein and reduced the binding affinity to -7.0 kcal/mol.

**Table 3:**
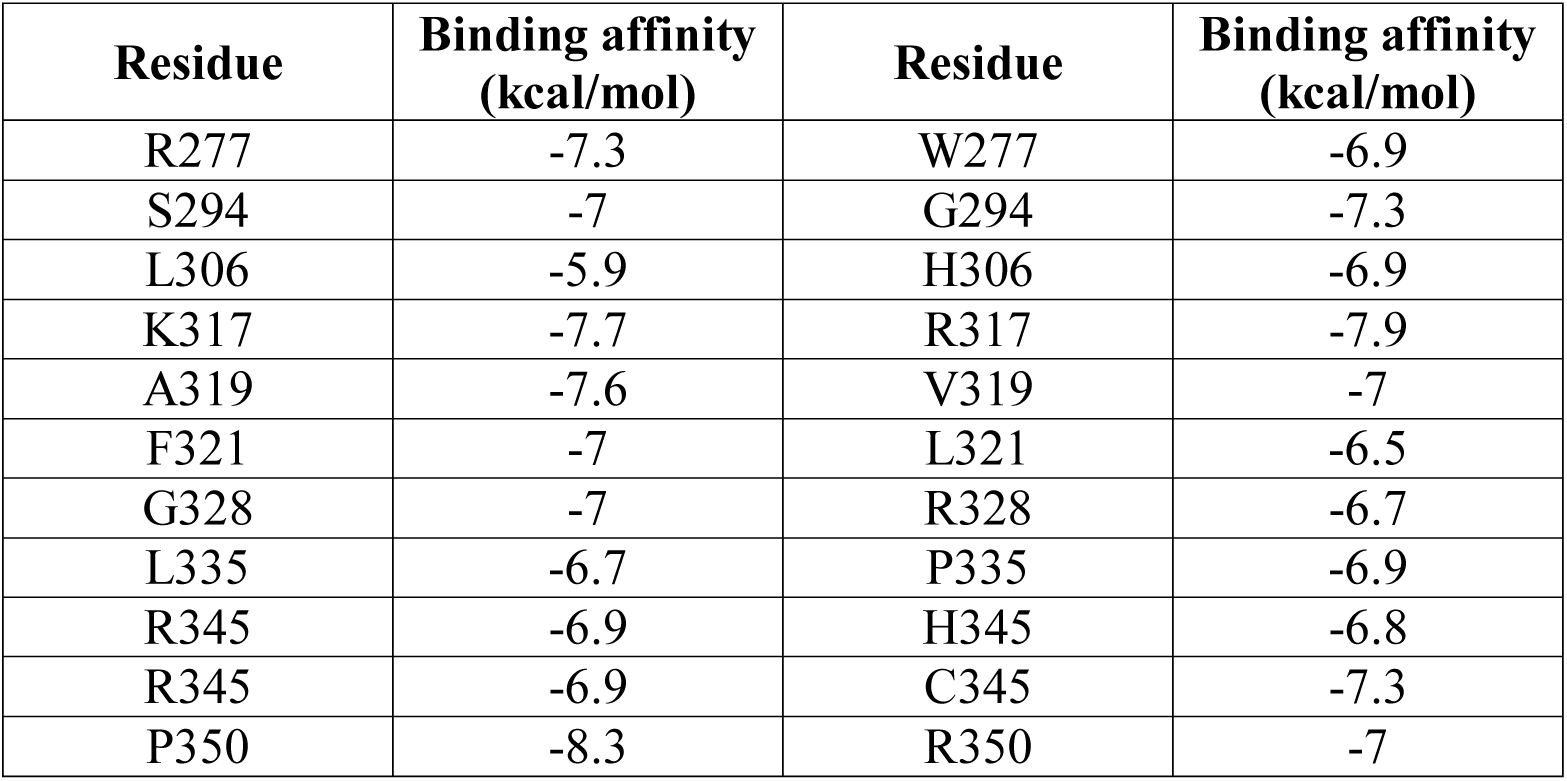
Docking results of RASSF5 with H-Ras.

| Residue | Binding affinity (kcal/mol) | Residue | Binding affinity (kcal/mol) |
| --- | --- | --- | --- |
| R277 | -7.3 | W277 | -6.9 |
| S294 | -7 | G294 | -7.3 |
| L306 | -5.9 | H306 | -6.9 |
| K317 | -7.7 | R317 | -7.9 |
| A319 | -7.6 | V319 | -7 |
| F321 | -7 | L321 | -6.5 |
| G328 | -7 | R328 | -6.7 |
| L335 | -6.7 | P335 | -6.9 |
| R345 | -6.9 | H345 | -6.8 |
| R345 | -6.9 | C345 | -7.3 |
| P350 | -8.3 | R350 | -7 |

**Figure 6:**
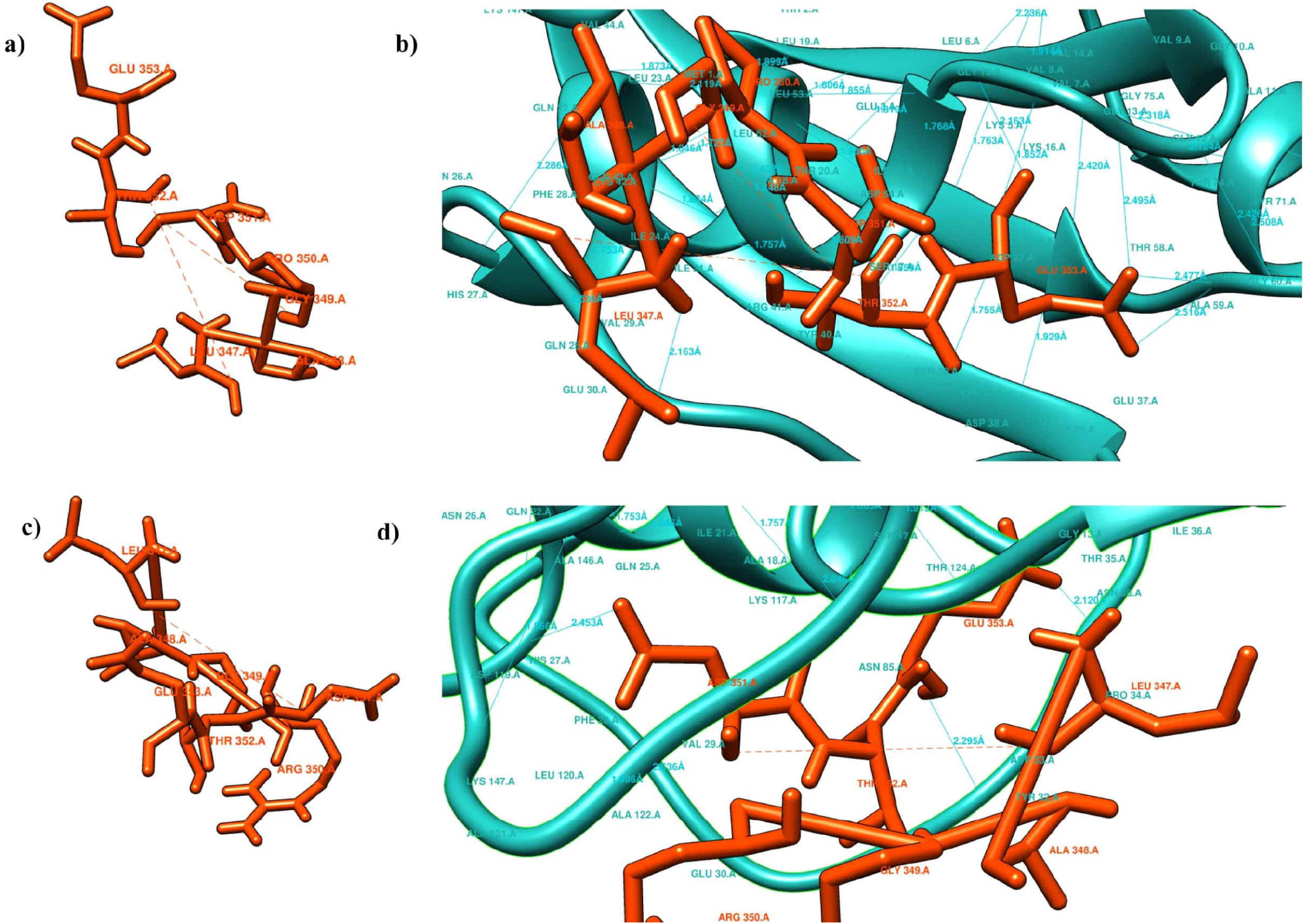
Molecular docking analysis of wild type and mutant peptide of P350R with H-Ras protein. **a)** Native peptide sequence (P350). **b)** Superimposed image of native peptide sequence and H-Ras protein. E353 forms two H-bonds with Gly60, one H-bond with G13 and another with G15. L347 forms one H-bond with E31. **c)** Mutant peptide sequence (R350). **d)** Superimposed image of mutant peptide sequence with H-Ras protein. Three new H-bonds are formed with the H-Ras protein where G353 forms two H-bonds with T35, and D33. Moreover, D351 forms single H-bond with K147.

### Prediction of post translational modification site

The GPS-MSP 1.0 tool predicted R277 as methylation site in RASSF5 protein with 90% specificity. We used NetPhos 3.1 and GPS 3.0 servers to predict the phosphorylation sites (**Figure 7**). Both tools predicted that wild type residues S294, S373 and mutant residues T379, Y154 would undergo phosphorylation (**Table S7)**. Moreover, UbPred tool predicted a total of eight (8) Lysine residues for ubiquitinylation but none was present in our highly risk nsSNPs. For further analysis of the 20 deleterious nsSNPs on PTM process, ModPred server was used to detect the proteolytic cleavage, ADP ribosylation, disulfide linkage, O-linked glycosylation, amidation sites in RASSF5. It predicted PTM sites in both wild and mutant residues of C137R, S260F, K317R, and R345H (**Table S8**).

**Figure 7:**
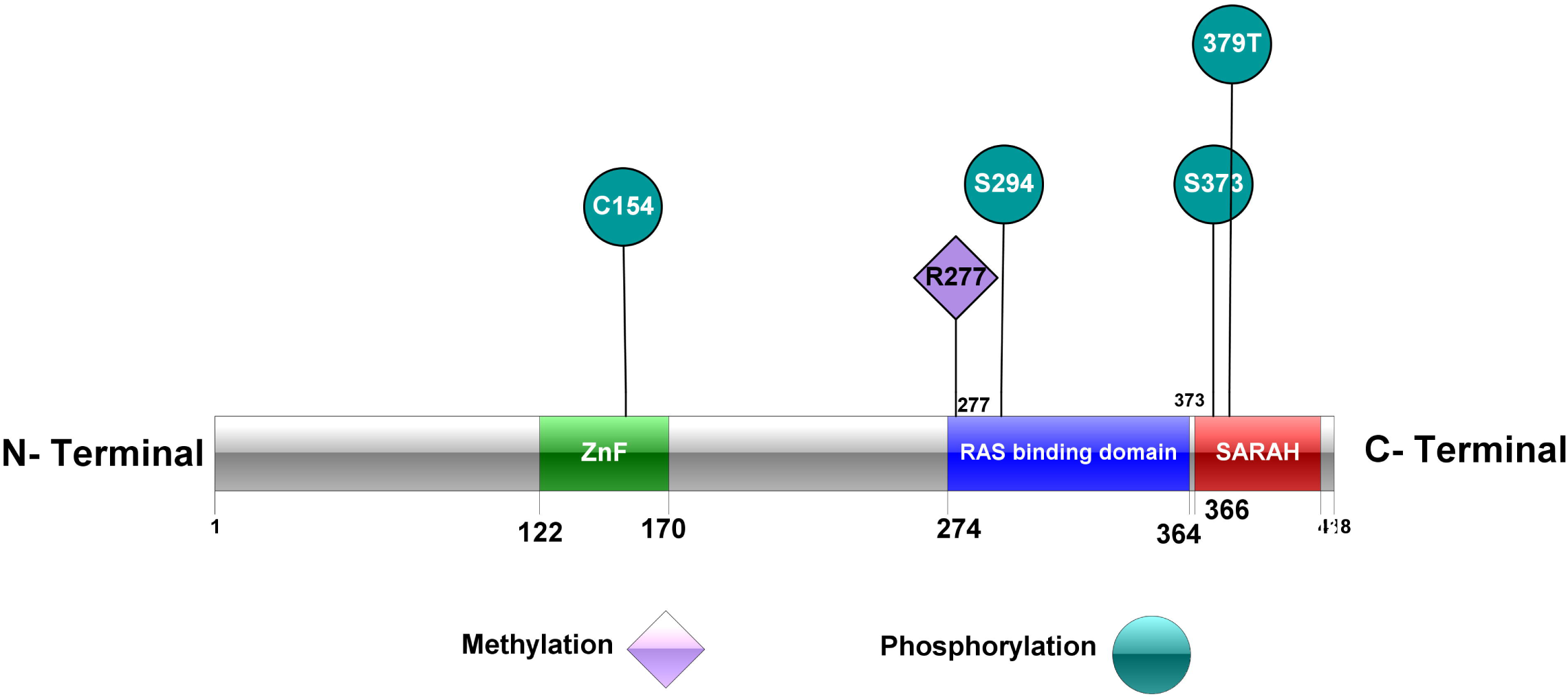
Post Translational Modification (Methylation and Phosphorylation) sites predicted by GPS-MSP 1.0, NetPhos 3.1 and GPS 3.0 (Using IBS software).

### Protein interacting network analysis

Cytoscape (v3.7.2) was used to build up the protein interaction network of RASSF5 protein and it predicted that RASSF5 protein is functionally associated with 18 proteins among which, HRAS, KRAS, NRAS, RRAS, RALGDS, RAP1A, RRAS2, RAP1B, MRAS, RAP1GAP, MLLT4, RAPGEF4 form one cluster and RASSF1, STK3, STK4, SAV1, MOB1A, MOB1B form another cluster. RASSF5 protein acts as a hub of these two clusters of proteins (**Figure 8**). If any change occurs in this RASSF5 protein, it may affect the overall protein network interaction among all these 18 proteins. Cytoscape also detected degree, average shortest path length, betweenness centrality, closeness centrality, neighborhood connectivity of all 18 proteins interacting with RASSF5 protein (**Table S9, S10**).

**Figure 8:**
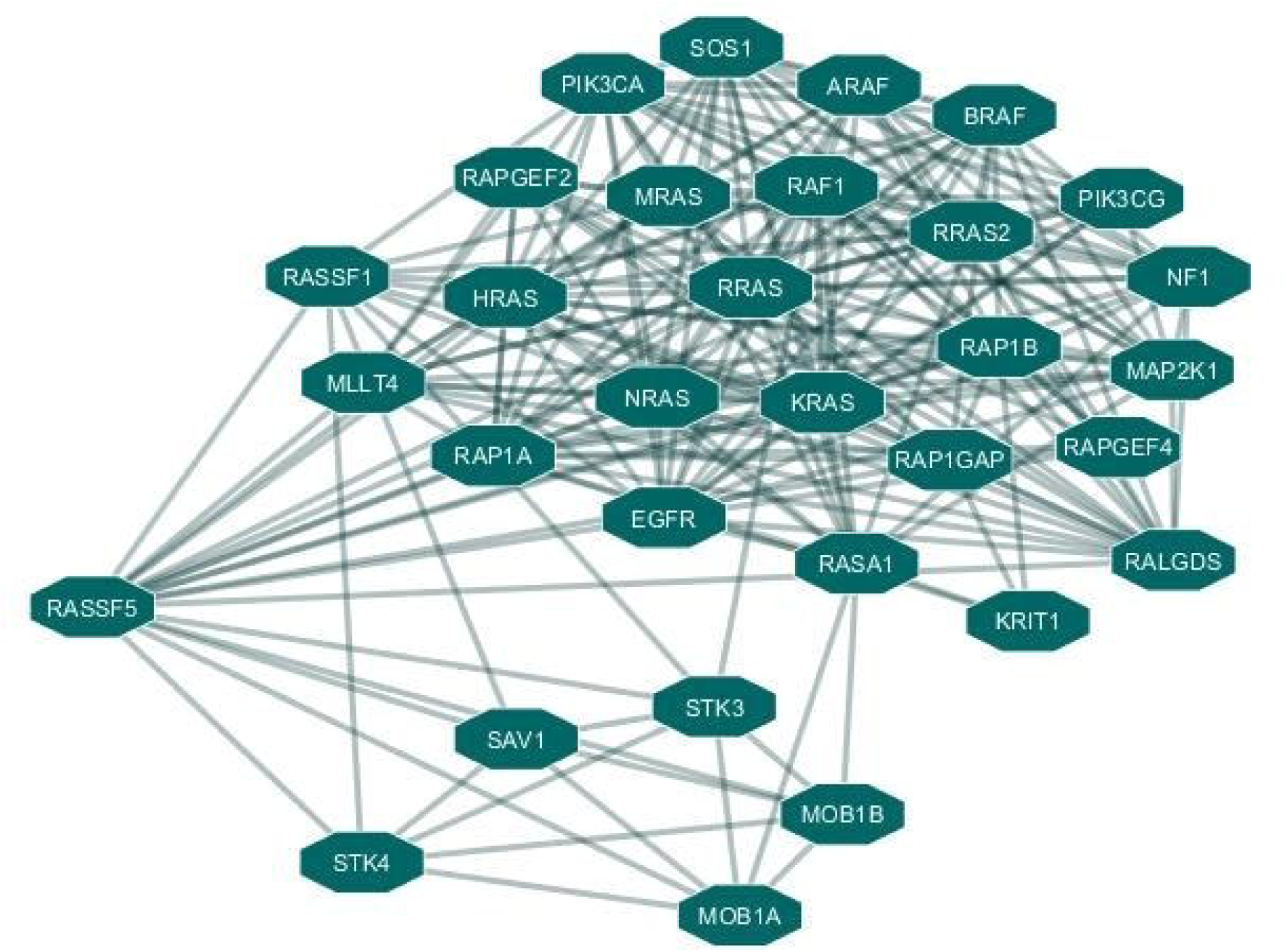
Protein interaction network of RASSF5 protein

## Discussion

Ras association domain-containing protein 5 (RASSF5), through their interaction with active GTP -bound Ras, acts as tumor suppressor in various human malignant tumors^3,29-33^. Inactivation of RASSF5 has been found in several human cancers, such as lung cancer^5^, colorectal cancer ^5^, gastric cancer^34^ etc. Changes in the structural conformation of RASSF5 protein during bio-molecular interactions play a vital role in executing its function^35^ but nsSNPs may cause aberrant conformations, which may lead to inactivation of its tumor suppressive properties^36,37^. Therefore, it is necessary to find out the effects of deleterious nsSNPs of RASSF5 and thereby, their association with various diseases. Here, we attempted computational analysis to find out the most deleterious nsSNPs and their effects on the structure and function of RASSF5 protein.

Among 495 nsSNPs found in NCBI database, we finally screened out 25 significantly deleterious nsSNPs using six *in silico* SNP prediction tools: SIFT, PANTHER, Polyphen-2, SNP&GO, PROVEAN, and PredictSNP. These 25 deleterious nsSNPs were selected on the basis of prediction scores generated by these six tools. We further used InterPro tool to determine the location of these nsSNPs on different domains of RASSF5. It revealed 20 nsSNPs in three different domains of the protein, where two nsSNPs were positioned in the Zinc finger domain that interacts with Ras association (RA) domain^38^, eleven (11) nsSNPs were located in the RA domain^3^, which serves as the binding site for active GTP-bound Ras, and the rest seven (7) nsSNPs were present in the SARAH domain that is crucial for cell apoptosis^39^.

Protein stability is essential for the structural and functional activity of a protein^40^. By using the I-Mutant tool, we obtained the deleterious nsSNPs that may affect the stability of the RASSF5 protein. Therefore, we focused on the consequences of the aforementioned 20 deleterious nsSNPs on the RASSF5 protein stability. Among these 20 nsSNPs, 17 nsSNPs decreased the stability of the protein. Protein stability governs the conformational structure of the protein and thus determines the function. Any alteration in protein stability may cause misfolding, degradation or aberrant conglomeration of proteins^41^. Moreover, evolutionary conservancy in the protein sequence is vital to determine whether a mutation has any negative effect on the host. Using the ConSurf web server, we found that highly deleterious nsSNPs having high conservation scores were located in the highly conserved regions, therefore, increases the risk of tumorigenesis by inactivating RASSF5.

The investigation continues with the analysis of the structural consequence of these deleterious nsSNPs using Phyre2 homology modeling tool. Using the three templates, we generated the wild type and mutant protein models for three domains of RASSF5 protein. Moreover, we used Tm Align tool that predicted RMSD and TM score of these wildtype and mutant protein models, which were generated by Phyre2. Higher RMSD and lower TM score signify higher deviation of mutant protein structure from native ones^25^. On the basis of these criteria, we finally selected 11 deleterious nsSNPs positioned in the RA domain. Remaining 9 nsSNPs show lower RMSD and higher TM score that indicates less deviation of mutant protein structure from wild type protein. I-TASSER provided more accurate structural and functional information of wild type and mutant protein models with C-score and Swiss Model provided the QMEN, solvation, torsion scores for our targeted 11 nsSNPs.

Project Hope server predicted that these 11 highly risky nsSNPs negatively affect the structure of the protein among which, 7 nsSNPs were structural and 4 nsSNPs were functional residues according to Consurf. Furthermore, Missense 3D tool predicted the consequences of the 7 structural nsSNPs and showed that 5 nsSNPs deleteriously affect the structural conformation of RASSF5 protein. Later on, we analyzed the energy minimization state of these 7 structural and 4 functional residues using Swiss PDB viewer, as the lowest energy conformation of the modeled protein is closer to the experimental structure^42^. The energy minimization score of wild type RASSF5 protein (−17442.365 kJ/mol) is significantly deviated in P350R (−16379.468 kJ/mol), F321L (−16381.121 kJ/mol), and R277W (−16448.330 kJ/mol). It was found that P350 forms three H-bonds with L347 (3.21 Å), E353 (3.26 Å), and V354 (2.92 Å) whereas, the mutant R350 clashes with V354. Besides this, variation in the energy minimization score compared to the wild type protein was observed in A319V, L335P, R345C, and R345H mutants as well.

Docking analysis confirmed that out of the 11 nsSNPs, four variants (P350R, A319V, F321L, and R277) significantly reduced the binding affinity with H-Ras protein compared to the wild type residues. The most remarkable change in binding affinity was observed in P350R where noticeable loss of H-bond interactions was found in the binding pocket. In the docking complex, Glu353 formed H-bonds with G60, G13, and G15 with distances of 2.477 Å, 2.516 Å, 2.495 Å, and 2.163 Å respectively and L347 formed H-bond with E31 with a distance of 2.163 Å. These H-bonds were disrupted when Arginine replaced Proline at 350th position, which altered the binding affinity. In short, molecular docking analysis revealed that the aforementioned variants can significantly affect the functional activity of RASSF5 protein.

GPS-MSP 1.0 predicted methylation site for R277 whereas, the variant W277 did not undergo methylation. Wild type residues (S294, S373) and mutant residues (T379, Y154) were the phosphorylation sites predicted by both NetPhos 3.1 and GPS 3.0 tools.

The protein interaction network of RASSF5 was determined based on degree, average shortest path length, betweenness centrality & closeness centrality using the Cytoscape tool. With the shortest path length as well as higher closeness & betweenness centrality, the result exhibited strong network interaction of RASSF5 with HRAS, KRAS, NRAS, RRAS, RALGDS, involved in the regulation of the signal transduction pathways, essential for cell growth, differentiation, migration, and apoptosis^43^. RAS bound with RASSF5, activate MST1/2^35,44^ to form STK3/MST2 and STK4/MST1 complexes in the hippo signaling pathway that plays a crucial role in tumor suppression by limiting proliferation and stimulating apoptosis where SAV1, MOB1A/B act as effector molecules^45,46^. Due to non-synonymous mutations in RASSF5, the signaling cascade might be disrupted, which in turn, may lead to tumorigenesis.

Ras association domain (RA) is required for the interaction between Ras family members and all Ras effectors. RASSF5 forms complex with active Ras through the RA domain, which thereby, activates MST1/2^32,39^. Activation of MST1/2 facilitates degradation of YAP1, whose over expression causes cancer^35^. If any deleterious mutation happens to occur in this RA domain, the overall tumor suppressive activity of RASSF5 will be significantly reduced. No such experimental study was documented till date for the protein RASSF5. So, this study will provide a substantial framework for the identification of functional SNPs.

## Methods

### Retrieving nsSNPs

From the National Center for Biotechnology Information (NCBI) dbSNP database (https://www.ncbi.nlm.nih.gov/projects/SNP), we retrieved the information of SNPs (SNP ID, protein accession number, location, residue alteration, and worldwide minor allele frequency (MAF)^47^. We filtered all the 495 nsSNPs for screening.

### Identifying the most deleterious SNPs

We recruited six distinct bioinformatics tools namely, **SIFT** (Sorting Intolerant From Tolerant)^48,49^ (http://sift.jcvi.org/www/SIFT_seq_submit2.html), **PANTHER** (Protein Analysis Through Evolutionary Relationship)^50^ (http://www.pantherdb.org/tools), **PolyPhen-2** (Polymorphism Phenotyping v2)^51^ (http://genetics.bwh.harvard.edu/pph2/), and **SNPs&GO**^52^ (http://snps.biofold.org/snps-and-go/snps-and-go.html), **PROVEAN** (Protein Variation Effect Analyzer)^53^ (http://provean.jcvi.org/index.php), and **PredictSNP**^54^ (http://loschmidt.chemi.muni.cz/predictsnp) to predict functional effects of the nsSNPs collected from dbSNP database. This ensured the accuracy and stringency of the results and we considered those SNPs as high-risk, which were anticipated as deleterious by all the six programs.

SIFT server uses sequence homology to interpret the effect of amino acid substitution to identify the tolerated and deleterious SNPs. Substitution of amino acid at a particular position with a probability <0.05 is considered as deleterious and intolerant, whereas, probability ≥ 0.05 is predicted as tolerant^29^. PANTHER program classifies the proteins on the basis of evolutionary relationship, molecular functions and their interactions with other proteins. It analyzes substitution based on position specific evolutionary conservation scores, which is calculated from alignment of various proteins that are evolutionarily related^50^. PolyPhen-2 predicts the functional effect of amino acid substitutions on the protein structure and functions based on sequence based characterization^51^. SNPs&GO server estimates the human disease related mutations based on support vector machines (SVM)^52^. PROVEAN, a web server that uses sequence homology to predict the functional effect of an amino acid substitution. The cutoff value of PROVEAN is set at -2.5. Substitutions of amino acid that exceed the cut off value were considered as deleterious. PredictSNP tool is a consensus SNP classifier, developed by exploiting six prediction programs (MAPP, PhD-SNP, PolyPhen-1, PolyPhen-2, SIFT and SNAP) to predict disease related mutations.

### Identification of nsSNPs on the domains of RASSF5

To identify the location of nsSNPs on the conserved domains of RASSF5 we used **InterPro**^9^ (http://www.ebi.ac.uk/interpro/) tool, which can recognize motifs, and domains of a protein and thereby identify the functional characterization of a protein by using the database consisting of protein families, domains and functional sites^55^.

### Analyzing the effect of the nsSNPs on protein stability

While engineering a protein, it is important to observe the extent to which a mutation affects the structure and stability of the protein. Thus, to serve this purpose, we used **I-Mutant 2.0**^56^ (http://folding.biofold.org/i-mutant/i-mutant2.0.html), which is a web server based on a support vector machine that predicts the stability of a protein after being mutated. This program utilizes the ProTherm-derived dataset, which is the largest database of experimental data on protein mutations^57^. We submitted the input data of 20 nsSNPs of RASSF5 in FASTA format.

### Analyzing protein evolutionary conservation

To identify the evolutionary conservation of the amino acids in the protein sequence we employed **ConSurf**^58^ (https://consurf.tau.ac.il) that performs its function by analyzing the phylogenetic relationships between homologous sequences^59,60^. We considered those nsSNPs of RASSF5 that were found to be conserved for further analyses.

### Prediction of structural effect of nsSNPs on human RASSF5 protein

To understand the effect of the nsSNPs on the structure of the protein we recruited **HOPE**^61^ (https://www3.cmbi.umcn.nl/hope). HOPE is a web-server that identifies the structural effects of the point mutations in a protein sequence. We used **Q8WWW0** (UniProt-Accession Code of RASSF5) and the 20 SNPs individually as the input.

Moreover, we also used **Missense 3D**^62^ (http://www.sbg.bio.ic.ac.uk/~missense3d/) and **Swiss-PDB Viewer**^63,64^ (v4.1.0) (https://spdbv.vital-it.ch/) to ensure the accuracy and stringency of our result. Missense 3D predicts the structural changes introduced by an amino acid substitution, whereas, Swiss-PDB Viewer is a stand-alone software that can generate mutated models of the proteins for the corresponding amino acid substitutions. The “Mutation Tool” of Swiss-PDB Viewer allows browsing through a rotamer library of the mutated models and the “best” rotamer of the mutated model can be selected. Then we performed energy minimization for the three-dimensional structures of the mutated models by utilizing **Nomad-Ref** server, which utilizes Gromacs as the ideal force field for energy minimization purpose^41^.

### Visualization of the effects of nsSNPs through 3D protein modeling

To create the 3D models of the mutated protein we used three different online tools: **SWISS-MODEL**^65^ (https://swissmodel.expasy.org/), **Phyre2**^66^ (http://www.sbg.bio.ic.ac.uk/phyre2), and **I-TASSER**^67-69^ (https://zhanglab.ccmb.med.umich.edu/I-TASSER/). As there was no crystal structure of RASSF5 available, we utilized Phyre2 and I-TASSER to predict the 3D structure and function of this protein. I-TAASER, especially, constructs full-length protein structures by splicing continuous threading alignments and then generates the structure by utilizing replica exchanged Monte Carlo simulations^67-69^. We used Ras domain-containing family protein 5 (3ddc.1.B), Ras association (RalGDS/AF-6) domain family 5 (1rfh.1.A), Ras association domain-containing protein 5 (4lgd.H) as models for this purpose. The templates were selected based on the score of confidence and sequence identity, generated by Phyre2 modeling tool (**Table S11**). These two factors are crucial in determining the probable accuracy of the model. Confidence denotes the probability of true homology based on the similarity between query sequence and template. Percentage of sequence identity between query and template needs to be above 30-40% for high accuracy model. After that we used **TM-align**^70^ (https://zhanglab.ccmb.med.umich.edu/TM-align/) to compare between native and mutated protein structures, which calculates template modeling-score (TM-score) and root mean square deviation (RMSD) and then superposes the structures. TM-score provides the result in 0 and 1, where 1 indicates perfect match between the structures. On the other hand, greater RMSD denotes greater variation between the native and the mutated structures. Finally, SWISS-MODEL was exploited, which is an automated protein structure homology-modeling server that uses the crystal structure of comparable protein as a model to assess native and mutant structures with various parameters.

### Molecular docking

To determine the effect of deleterious point mutations over the binding affinity of RASSF5, we performed molecular docking using the **PyRx** virtual screening tool^71^ (https://pyrx.sourceforge.io/). We prepared suitable target protein from crystal structure complex of RASSF5 with active Ras protein found in PDB using **Discovery Studio**^72^ (v4.5). The peptide sequences from mutated RASSF5 protein containing nsSNPs are used as the ligands for the docking procedure. The PDB format of these input ligands were converted into pdbqt format using **Python Molecular Viewer** with Autodock instruments, **AutodockVina**^73,74^. We maximized the grid box size along the axes X=38.709, Y=43.839, Z=43.929 respectively. The docking result and the binding interaction between ligand and receptor proteins were visualized by **UCSF Chimera 1.13rc** tool^75^.

### Prediction of different post-translational modification sites

Post-translation modifications (PTM) influence a range of significant biological procedures including cell signaling, metabolic pathways etc. Putative PTM sites in the human RASSF5 protein were identified by using the web-server **ModPred**^76^ (www.modpred.org/). To predict potential sites which can accommodate phosphorylation, we utilized **NetPhos 3.1**^77^ (http://www.cbs.dtu.dk/services/NetPhos/) and **GPS 5.0**^78^ (http://gps.biocuckoo.cn/). NetPhos 3.1 server predicts the Serine, Threonine and Tyrosine phosphorylation sites in the protein by using ensembles of neural networks. Residues in the protein those have the score greater than 0.5 indicates phosphorylation. Similarly, higher score in the GPS 5.0 corresponds to higher probability of getting phosphorylated.

Apart from identifying the phosphorylation sites, we also predicted the potential sites for methylation by **GPS-MSP 1.0**^78^ (http://msp.biocuckoo.org/), ubiquitylation by **UbPred**^16^ (www.ubpred.org) and **BDM-PUB** (www.bdmpub.biocuckoo.org), and SUMOylation by using the tool **GPS-SUMO**^79,80^ (http://sumosp.biocuckoo.org/).

### Prediction of molecular interaction networks

Maintaining the interactions among proteins is very crucial to conserve homeostasis of the system. **Cytoscape**^81^ (v3.7.2) is a freely available Java based software program that was used to visualize molecular interaction networks of human RASSF5 protein with other associated proteins. Visualization of interacting networks of several types of a protein can be done only if the data is introduced in Simple Interaction Format (SIF) or GML format^82^.

## Conclusion

RASSF5 protein plays a vital role as tumor suppressor. One of the functional domains of this protein is the Ras-association (RA) domain that interacts with GTP-bound H-Ras to execute its tumor suppressive role. Thus, the structural conformation of this domain is very crucial for exerting its functional role. This *in silico* analysis of the functional SNPs of *RASSF5* provides significant insight into the deleterious effects that the nsSNPs might cause to the protein. This is the first study that predicts the impacts of nsSNPs on the structure and function of RASSF5 protein. Our study reports that four nsSNPs (P350R, A319V, F321L, and R277W) in the RA domain reduce the binding affinity with H-Ras, where P350R shows the most significant reduction. As this protein has been found to be involved in various tumors, our findings will serve as important guide in the studies of potential diagnostic and therapeutic interventions which require experimental mutational validation and clinical trial-based studies on a large population.

## Data availability

All data generated or analyzed during this study are included in this article and its Supplementary Information files.

## Acknowledgements

The authors acknowledge the Department of Biotechnology & Genetic Engineering, Noakhali Science & Technology University and the Department of Biochemistry & Biotechnology, University of Barishal for providing the research work supports.

## Author Contributions

MSH and MSI conceived the project and designed the study. MSH, ASR and MSI performed the analyses. MSH and MSI wrote the manuscript. All authors reviewed the manuscript.

## Additional Information

**Competing Interests Statement:** The authors declare no competing interests.

## Supplementary Information

Supplementary information for this study may be found online in the Supplementary Information section.

