## Supplementary material for "*In Silico* Analysis Predicting Effects of Deleterious SNPs of Human *RASSF5* Gene on its Structure and Functions": Table S1-Table S10

**Table S1:** High risk nsSNPs identified by four *in silico* programs

| **S.N.** | **SNP Id** | **AA substitution** | **SIFT Prediction** | **Polyphen Prediction** | **Panther Prediction** | **SNPs&GO Prediction** |
| --- | --- | --- | --- | --- | --- | --- |
| 1. | rs1344063691 | C137R | Deleterious | Probably Damaging | probably damaging | Disease |
| 2. | rs781984025 | C154Y | Deleterious | Probably Damaging | probably damaging | Disease |
| 3. | rs199649479 | G234S | Deleterious | Probably Damaging | probably damaging | Disease |
| 4. | rs782389172 | G238D | Deleterious | Probably Damaging | probably damaging | Disease |
| 5. | rs369854759 | R247W | Deleterious | Probably Damaging | probably damaging | Disease |
| 6. | rs545894272 | R248Q | Deleterious | Probably Damaging | probably damaging | Disease |
| 7. | rs539554161 | S260F | Deleterious | Probably Damaging | possibly damaging | Disease |
| 8. | rs4845112 | R277W | Deleterious | Probably Damaging | probably damaging | Disease |
| 9. | rs782500219 | S294G | Deleterious | Probably Damaging | probably damaging | Disease |
| 10. | rs782662751 | L306H | Deleterious | Probably Damaging | probably damaging | Disease |
| 11. | rs200663011 | K317R | Deleterious | Probably Damaging | probably damaging | Disease |
| 12. | rs782205958 | A319V | Deleterious | Probably Damaging | probably damaging | Disease |
| 13. | rs1054565713 | F321L | Deleterious | Probably Damaging | Probably damaging | Disease |
| 14. | rs144913020 | G328R | Deleterious | Probably Damaging | probably damaging | Disease |
| 15. | rs746023356 | L335P | Deleterious | Probably Damaging | probably damaging | Disease |
| 16. | rs782213884 | R345H | Deleterious | Probably Damaging | probably damaging | Disease |
| 17. | rs782615062 | R345C | Deleterious | Probably Damaging | probably damaging | Disease |
| 18. | rs1171545733 | P350R | Deleterious | Probably Damaging | probably damaging | Disease |
| 19. | rs563800378 | V367A | Deleterious | Probably Damaging | probably damaging | Disease |
| 20. | rs782710434 | F372L | Deleterious | Probably Damaging | probably damaging | Disease |
| 21. | rs374894198 | S373C | Deleterious | Probably Damaging | probably damaging | Disease |
| 22. | rs150973687 | P375R | Deleterious | Probably Damaging | probably damaging | Disease |
| 23. | rs368719396 | N379T | Deleterious | Probably Damaging | probably damaging | Disease |
| 24. | rs782097670 | N379H | Deleterious | Probably Damaging | probably damaging | Disease |
| 25. | rs782502649 | E388Q | Deleterious | Probably Damaging | probably damaging | Disease |

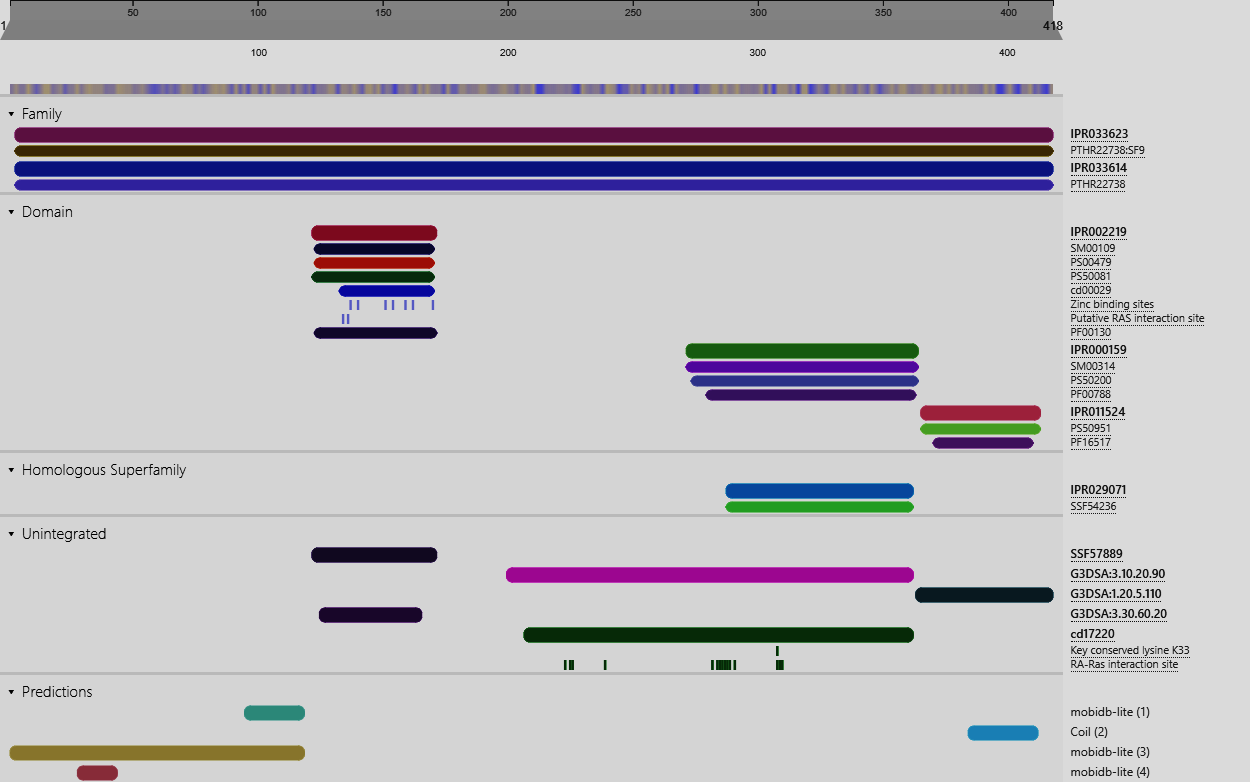

**Figure S1**: Domain identification of RASSF5 protein using InterPRO server. Here, IPR033623 indicates the RASSF5 protein (3-418aa), [IPR002219](https://www.ebi.ac.uk/interpro/entry/InterPro/IPR002219/) indicates Zinc finger domain (122-170), [IPR000159](https://www.ebi.ac.uk/interpro/entry/InterPro/IPR000159/) indicates Ras-association domain (274-364), and [IPR011524](https://www.ebi.ac.uk/interpro/entry/InterPro/IPR011524/) indicates SARAH domain (366-413).

**Table S2:** Effect of nsSNPs on protein stability predicted by I-MUTANT 2.0

| **Serial No.** | **SNP Id** | **AA substitution** | **I-Mutant** | **RI** | **DDG** |
| --- | --- | --- | --- | --- | --- |
| 1. | rs1344063691 | C137R | Decrease | 4 | -0.9 |
| 2. | rs781984025 | C154Y | Decrease | 3 | 0.82 |
| 3. | rs4845112 | R277W | Decrease | 6 | -0.83 |
| 4. | rs782500219 | S294G | Decrease | 7 | -1.6 |
| 5. | rs782662751 | L306H | Decrease | 9 | -2.43 |
| 6. | rs200663011 | K317R | Increase | 3 | -0.05 |
| 7. | rs782205958 | A319V | Decrease | 3 | -1.71 |
| 8. | rs1054565713 | F321L | Decrease | 8 | -2.56 |
| 9. | rs144913020 | G328R | Decrease | 7 | -1.36 |
| 10. | rs746023356 | L335P | Decrease | 5 | -1.46 |
| 11. | rs782213884 | R345H | Decrease | 9 | -1.37 |
| 12. | rs782615062 | R345C | Decrease | 6 | -1.14 |
| 13. | rs1171545733 | P350R | Decrease | 7 | -1.59 |
| 14. | rs563800378 | V367A | Decrease | 9 | -0.34 |
| 15. | rs782710434 | F372L | Decrease | 6 | -1.37 |
| 16. | rs374894198 | S373C | Increase | 0 | -0.89 |
| 17. | rs150973687 | P375R | Decrease | 7 | -0.72 |
| 18. | rs368719396 | N379T | Increase | 0 | -0.53 |
| 19. | rs782097670 | N379H | Decrease | 7 | -1.8 |
| 20. | rs782502649 | E388Q | Decrease | 1 | 0.06 |

**Table S3:** Evolutionary conservancy of amino acids in RASSF5 analyzed by Consurf

| **SNP Id** | **Residue and Position** | **Conservation Score** | **Prediction** |
| --- | --- | --- | --- |
| rs1344063691 | C137 | 8 | conserved and Buried |
| rs781984025 | C154 | 9 | Highly conserved and Buried (S) |
| rs199649479 | G234 | 8 | Highly Conserved and Exposed (F) |
| rs782389172 | G238 | 9 | Highly conserved and Buried (S) |
| rs369854759 | R247 | 6 | Average conserved and Exposed |
| rs545894272 | R248 | 9 | Highly Conserved and Exposed (F) |
| rs539554161 | S260 | 4 | Average conserved and Exposed |
| rs4845112 | R277 | 8 | Highly Conserved and Exposed (F) |
| rs782500219 | S294 | 9 | Highly conserved and Buried (S) |
| rs782662751 | L306 | 9 | Highly conserved and Buried (S) |
| rs200663011 | K317 | 9 | Highly Conserved and Exposed (F) |
| rs782205958 | A319 | 9 | Highly conserved and Buried (S) |
| rs1054565713 | F321 | 8 | Conserved and Buried |
| rs144913020 | G328 | 6 | Average conserved and Exposed |
| rs746023356 | L335 | 9 | Highly conserved and Buried (S) |
| rs782213884 | R345 | 9 | Highly conserved and Buried (S) |
| rs782615062 | R345 |  |  |
| rs1171545733 | P350 | 9 | Highly Conserved and Exposed (F) |
| rs563800378 | V367 | 9 | Highly conserved and Buried (S) |
| rs782710434 | F372 | 9 | Highly conserved and Buried (S) |
| rs374894198 | S373 | 9 | Highly conserved and Buried (S) |
| rs150973687 | P375 | 9 | Highly Conserved and Exposed (F) |
| rs368719396 | N379 | 9 | Highly Conserved and Exposed (F) |
| rs782097670 | N379 |  |  |
| rs782502649 | E388 | 9 | Highly Conserved and Exposed (F) |

**Table S4:** TM-align predictions for nsSNPs in RASSF5

| Serial No. | AA substitution | Domain | Tm Score | RMSD |
| --- | --- | --- | --- | --- |
| 1. | C137R | Zinc finger domain | 0.95234 | 0.72 |
| 2. | C154Y |  | 0.97681 | 0.46 |
| 3. | R277W | Ras-association domain | 0.85619 | 1.95 |
| 4. | S294G |  | 0.84095 | 1.08 |
| 5. | L306H |  | 0.86561 | 1.76 |
| 6. | K317R |  | 0.85966 | 2.06 |
| 7. | A319V |  | 0.87086 | 1.39 |
| 8. | F321L |  | 0.90398 | 1.54 |
| 9. | G328R |  | 0.86187 | 1.64 |
| 10. | L335P |  | 0.87452 | 1.64 |
| 11. | R345H |  | 0.87366 | 2.19 |
| 12. | R345C |  | 0.89856 | 1.6 |
| 13. | P350R |  | 0.85379 | 1.43 |
| 14. | V367A | SARAH domain | 0.98902 | 0.19 |
| 15. | F372L |  | 0.99788 | 0.08 |
| 16. | S373C |  | 0.88636 | 0.72 |
| 17. | P375R |  | 0.98342 | 0.24 |
| 18. | N379T |  | 0.99755 | 0.09 |
| 19. | N379H |  | 0.99707 | 0.1 |
| 20. | E388Q |  | 0.99187 | 0.17 |

**Table S5:** Comparative impacts of 11 nsSNPs on 3D structure of RASSF5 investigated by Swiss Model

| **SNPs** | **QMEN** | **Cb** | **All atom** | **Solvation** | **Torsion** | **Template** |
| --- | --- | --- | --- | --- | --- | --- |
| Wild Type | -1.27 | -1.06 | -0.32 | -0.16 | -1.03 | 3ddc.1.B |
| R277W | -1.8 | -1.18 | -0.58 | 0.34 | -1.72 | 3ddc.1.B |
| S294G | -1.39 | -1.06 | -0.3 | -0.16 | -1.14 | 3ddc.1.B |
| L306H | -1.32 | -0.9 | -0.37 | -0.27 | -1.07 | 3ddc.1.B |
| K317R | -1.64 | -0.89 | -0.38 | -0.31 | -1.39 | 3ddc.1.B |
| A319V | -1.21 | -0.94 | -0.39 | -0.22 | -0.96 | 3ddc.1.B |
| F321L | -1.41 | -1.2 | -0.38 | -0.3 | -1.08 | 3ddc.1.B |
| G328R | -1.49 | -1.19 | -0.32 | -0.04 | -1.27 | 3ddc.1.B |
| L335P | -1.19 | -0.87 | -0.48 | -0.18 | -0.96 | 3ddc.1.B |
| R345H | -1.26 | -0.89 | -0.39 | -0.17 | -1.04 | 3ddc.1.B |
| R345C | -1.37 | -0.96 | -0.35 | -0.2 | -1.13 | 3ddc.1.B |
| P350R | -1.42 | -0.96 | -0.39 | -0.14 | -1.2 | 3ddc.1.B |

**Table S6:** Structural effect of 11 nsSNPs over RASSF5 protein using Project Hope

| **Residue** | **Structure** | **Properties** |
| --- | --- | --- |
| **R277W** | 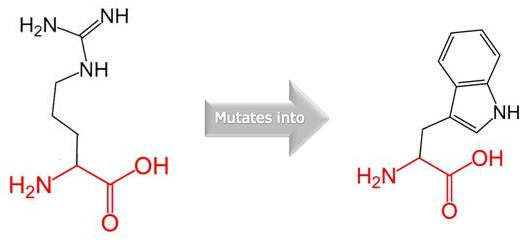 | - Mutant residue is bigger than the wild-type residue. - Wild-type residue charge was POSITIVE, the mutant residue charge is NEUTRAL. - Mutant residue is more hydrophobic than the wild-type residue. - The mutation is located within Ras-associating domain, which can disturb this domain and abolish its function. |
| **S294G** | 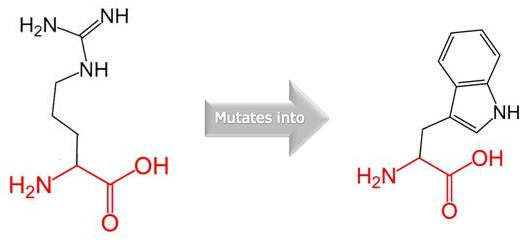 | - Mutant residue is smaller than the wild-type residue. - New residue is not in the correct position to make the same hydrogen bond as the original wild-type residue did. - Mutation introduces an amino acid with different properties, can disturb this domain and abolish its function. - Mutation introduces a glycine which are very flexible and can disturb the required rigidity of the protein at this position. |
| **L306H** | 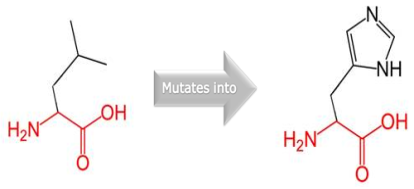 | - Mutant residue is bigger than the wild-type residue. - Wild-type residue is more hydrophobic than the mutant residue. - The mutation introduces an amino acid with different properties, which can disturb this domain and abolish its function. |
| **K317R** | 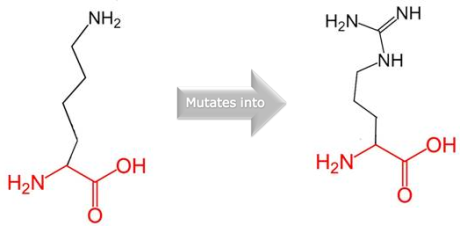 | - Mutant residue is bigger than the wild-type residue. - The mutation is located within Ras-associating domain, which can disturb this domain and abolish its function |
| **A319V** | 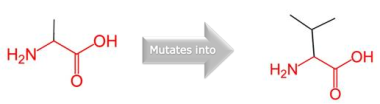 | - Mutant residue is bigger than the wild-type residue - The mutation is located within Ras-associating domain, which can disturb this domain and abolish its function |
| **F321L** | 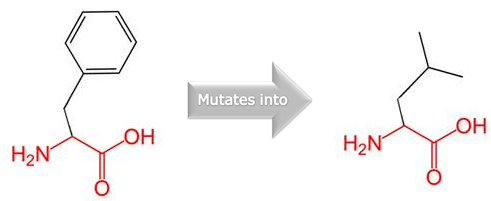 | - Mutant residue is smaller than the wild-type residue. - The mutation is located within Ras-associating domain, which can disturb this domain and abolish its function |
| **G328R** | 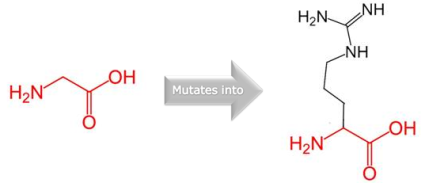 | - The mutation is located within Ras-associating domain, which can disturb this domain and abolish its function. - Wild-type residue is a glycine, mutation of this glycine can abolish flexibility function. |
| **L335P** | 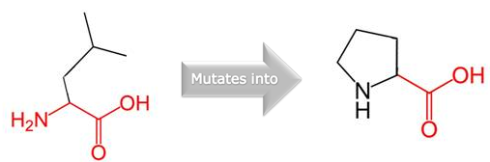 | - The mutant residue is smaller than the wild-type residue. - The mutation is located within Ras-associating domain, which can disturb this domain and abolish its function. |
| **R345H** | 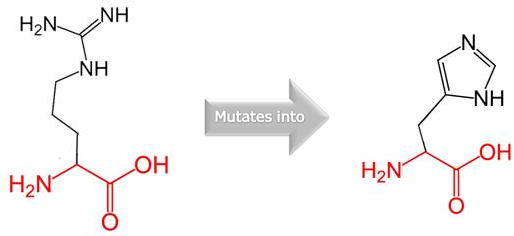 | - Mutant residue is smaller than the wild-type residue. - Wild-type residue charge was POSITIVE whereas, mutant residue is NEUTRAL. - The new residue is not in the correct position to make the same hydrogen bond as the wild-type residue did. - Wild-type residue forms a salt bridge at position 222, at position 351 - Difference in charge will disturb the ionic interaction. |
| **R345C** | 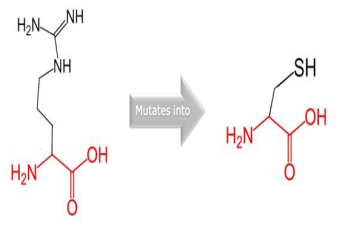 | - Mutant residue is smaller than the wild-type residue. - Wild-type residue charge was POSITIVE, the mutant residue charge is NEUTRAL. - Mutant residue is more hydrophobic than the wild-type residue. - The wild-type residue forms a hydrogen bond at position 355 - The new residue is not in the correct position to make the same hydrogen bond as the original wild-type residue did. - The difference in hydrophobicity will affect hydrogen bond formation. - The wild-type residue forms a salt bridge at position 222 and at position 351 - Difference in charge will disturb the ionic interaction made by wild-type residue. |
| **P350R** | 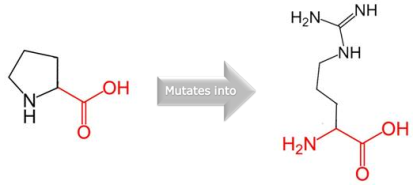 | - Mutant residue is bigger than the wild-type residue. - Wild-type residue charge was NEUTRAL whereas the mutant residue is POSITIVE. - Wild-type residue is more hydrophobic than the mutant residue - The mutation is located within Ras-associating domain, which can disturb this domain and abolish its function. - The wild-type residue is a proline. and therefore, induce a special backbone conformation which might be required at this position and mutation can disturb this special conformation |

**Table S7:** Structural effect of 11 nsSNPs over RASSF5 protein using Missense3D tool

| **AA substitution** | **PDB ID** | **Chain** | **Result Analysis** | **Detailed Analysis** |
| --- | --- | --- | --- | --- |
| S294G | 3ddc | B | No structural damage detected |  |
| L306H | 3ddc | B | - Buried hydropilic introduced - Buried charge introduced | - This substitution replaces a buried hydrophobic residue (LEU, RSA 1.2%) with a hydrophilic residue (HIS, RSA 1.0%). - This substitution replaces a buried uncharged residue (LEU, RSA 1.2%) with a charged residue (HIS). |
| A319V | 3ddc | B | - Buried / exposed switch | - This substitution results in a change between buried and exposed state of the target variant residue. ALA is buried (RSA 3.7%) and VAL is exposed (RSA 19.7%). |
| F321L | 3ddc | B | No structural damage detected |  |
| L335P | 3ddc | B | - Buried Pro introduced | - This substitution introduces a buried proline. |
| R345H | 3ddc | B | - Buried H-bond breakage - Buried / exposed switch | - This substitution disrupts all side-chain / side-chain H-bond(s) and/or side-chain / main-chain bond(s) H-bonds formed by a buried ARG residue (RSA 2.8%). - This substitution results in a change between buried and exposed state of the target variant residue. ARG is buried (RSA 2.8%) and HIS is exposed (RSA 12.5%). |
| R345C | 3ddc | B | - Cavity altered | - The substitution leads to the expansion of cavity volume by 98.064 Å^3 |

**Table S8:** Swiss PDB Viewer Result

| **SL** | **AA substitution** | **Presence of Clash / Hydrogen bond** | **Number of Rotamer** | **Total Energy after Energy Minimization**  **(I-TASSER)** |
| --- | --- | --- | --- | --- |
| 1. | Wild type |  |  | -17442.365 |
| 2. | R277W | Both | 7 | -16448.330 |
| 3. | S294G | Hydrogen bond | 1 | -17319.016 |
| 4. | L306H | Both | 6 | -17438.125 |
| 5. | K317R | Both | 28 | -17579.576 |
| 6. | A319V | Both | 3 | -17243.609 |
| 7. | F321L | BOTH | 4 | -16381.121 |
| 8. | G328R | Both | 28 | -16816.518 |
| 9. | L335P | BOTH | 2 | -17017.646 |
| 10. | R345H | BOTH | 6 | -17134.549 |
| 11. | R345C | Hydrogen bond | 3 | -17142.982 |
| 12. | P350R | Both | 28 | -16379.468 |

**Table S9 a):** Prediction of Phosphorylation sites in wild type RASSF5 using NETPHOS3.1

| **Netphos 3.1** | **Wild Type** | | | **Netphos 3.1** | **Mutant Residues** | | |
| --- | --- | --- | --- | --- | --- | --- | --- |
| **Serine (S)** | **Position** | **Score** | **Kinase** | **Serine (S)** | **Position** | **Score** | **Kinase** |
|  | 5 S | 0.544 | p38MAPK |  | 5 S | 0.544 | p38MAPK |
|  | 27 S | 0.544 | PKA |  | 27 S | 0.544 | PKA |
|  | 29 S | 0.746 | unsp |  | 29 S | 0.746 | unsp |
|  | 41 S | 0.462 | GSK3 |  | 41 S | 0.462 | GSK3 |
|  | 42 S | 0.766 | unsp |  | 42 S | 0.766 | unsp |
|  | 51 S | 0.454 | GSK3 |  | 51 S | 0.454 | GSK3 |
|  | 61 S | 0.983 | unsp |  | 61 S | 0.983 | unsp |
|  | 77 S | 0.796 | unsp |  | 77 S | 0.796 | unsp |
|  | 105 S | 0.977 | unsp |  | 105 S | 0.977 | unsp |
|  | 164 S | 0.723 | PKA |  | 164 S | 0.723 | PKA |
|  | 171 S | 0.894 | unsp |  | 171 S | 0.894 | unsp |
|  | 177 S | 0.868 | unsp |  | 177 S | 0.868 | unsp |
|  | 182 S | 0.998 | unsp |  | 177 S | 0.868 | unsp |
|  | 185 S | 0.528 | cdc2 |  | 182 S | 0.998 | unsp |
|  | 192 S | 0.634 | DNAPK |  | 185 S | 0.528 | cdc2 |
|  | 217 S | 0.498 | unsp |  | 192 S | 0.634 | DNAPK |
|  | 231 S | 0.527 | CKI |  | 217 S | 0.498 | unsp |
|  | 260 S | 0.997 | unsp |  | 231 S | 0.522 | PKA |
|  | 279 S | 0.992 | unsp |  | 234 S | 0.908 | unsp |
|  | 293 S | 0.536 | PKA |  | 279 S | 0.451 | CaM-II |
|  | 294 S | 0.448 | CaM-II |  | 293 S | 0.589 | PKA |
|  | 299 S | 0.450 | CaM-II |  | 356 S | 0.507 | CKII |
|  | 336 S | 0.561 | PKA |  | 299 S | 0.448 | CaM-II |
|  | 356 S | 0.515 | CKII |  | 336 S | 0.540 | PKA |
|  | 373 S | 0.579 | CKI |  | 413 S | 0.958 | unsp |
|  | 413 S | 0.958 | unsp |  | 492 S | 0.958 | unsp |
| **Threonine (T)** | 52 T | 0.491 | PKG | **Threonine (T)** | 52 T | 0.491 | PKG |
|  | 152 T | 0467 | cdc2 |  | 152 T | 0.498 | PKC |
|  | 157 T | 0.458 | cdc2 |  | 157 T | 0.475 | cdc2 |
|  | 186 T | 0.866 | unsp |  | 186 T | 0.866 | unsp |
|  | 188 T | 0.632 | PKC |  | 188 T | 0.632 | PKC |
|  | 190 T | 0.549 | PKC |  | 190 T | 0.549 | PKC |
|  | 202 T | 0.977 | unsp |  | 202 T | 0.977 | unsp |
|  | 207 T | 0.467 | GSK3 |  | 207 T | 0.467 | GSK3 |
|  | 220 T | 0.439 | GSK3 |  | 220 T | 0.439 | GSK3 |
|  | 235 T | 0.477 | DNAPK |  | 235 T | 0.446 | DNAPK |
|  | 237 T | 0.443 | GSK3 |  | 237 T | 0.462 | cdc2 |
|  | 251 T | 0.460 | CaM-II |  | 251 T | 0.436 | CaM-II |
|  | 273 T | 0.774 | PKC |  | 273 T | 0.657 | PKC |
|  | 274 T | 0.945 | unsp |  | 274 T | 0.915 | unsp |
|  | 278 T | 0.493 | PKG |  | 278 T | 0.440 | CaM-II |
|  | 295 T | 0.772 | PKC |  | 295 T | 0.787 | PKC |
|  | 296 T | 0.594 | unsp |  | 296 T | 0.752 | unsp |
|  | 297 T | 0.932 | unsp |  | 297 T | 0.956 | unsp |
|  | 352 T | 0.483 | unsp |  | 352 T | 0.912 | unsp |
|  | 364 T | 0.592 | CKII |  | 364 T | 0.606 | CKII |
|  | 382 T | 0.614 | CKII |  | 379 T | 0.464 | CaM-II |
| **Tyrosine (Y)** | 13 Y | 0.380 | INSR | **Tyrosine (Y)** | 382 T | 0.497 | CKII |
|  | 24 Y | 0.804 | unsp |  | 443 T | 0.606 | CKII |
|  | 218 Y | 0.486 | INSR |  | 461 T | 0.538 | CKII |
|  | 236 Y | 0.531 | unsp |  | 13 Y | 0.380 | INSR |
|  | 262 Y | 0.498 | unsp |  | 24 Y | 0.804 | unsp |
|  | 281 Y | 0.408 | INSR |  | 154 Y | 0.646 | unsp |
|  | 343 Y | 0.496 | unsp |  | 218 Y | 0.486 | INSR |
|  | 399 Y | 0.413 | INSR |  | 236 Y | 0.952 | unsp |

** Result of GPS 5.0 is provided in the Supplementary Files 2 and 3.

**Table S10:** Modpred result of all 20 deleterious nsSNPs

| **nsSNPs** | **Residue (wild type & mutant)** | **Modification** | **Score** | **Confidence** | **Remarks** |
| --- | --- | --- | --- | --- | --- |
| **C137R** | C137 | Disulfide linkage | 0.52 | Low | Novel prediction |
|  | R137 | Proteolytic cleavage | 0.53 | Low | Novel prediction |
| **C154Y** | C154 | Disulfide linkage | 0.63 | Low | Novel prediction |
|  | Y154 |  |  |  |  |
| **S294G** | S294 | O-linked glycosylation | 0.58 | Low | S294 |
|  | G294 |  |  |  |  |
| **K317R** | K317 | Acetylation | 0.73 | Medium | Novel prediction |
|  | K317 | Proteolytic cleavage | 0.57 | Low | Novel prediction |
|  | R317 | Proteolytic cleavage | 0.80 | Medium | Novel prediction |
| **L335P** | L335 |  |  |  |  |
|  | P335 | Proteolytic cleavage | 0.64 | Low | Novel prediction |
| **R345H** | R345 | ADP-ribosylation | 0.58 | Low | Novel prediction |
|  | R345 | Proteolytic cleavage | 0.71 | Medium | Novel prediction |
|  | H345 | Proteolytic cleavage | 0.6 | Low | Novel prediction |
| **P350R** | P350 |  |  |  |  |
|  | R350 | Proteolytic cleavage | 0.72 | Medium | Novel prediction |
| **F372L** | F372 | Proteolytic cleavage | 0.57 | Low | Novel prediction |
|  | L372 |  |  |  |  |
| **S373C** | S373 | Proteolytic cleavage | 0.51 | Low | Novel prediction |
|  | C373 |  |  |  |  |
| **P375R** | P375 |  |  |  |  |
|  | R375 | ADP-ribosylation | 0.7 | Medium | Novel prediction |
| **N379T/H** | N379 | Proteolytic cleavage | 0.56 | Low | Novel prediction |
|  | T/H379 |  |  |  |  |
| **E388Q** | E388 |  |  |  |  |
|  | Q388 | Pyrrolidone carboxylic acid | 0.57 | Medium | Novel prediction |
|  | Q388 | Amidation | 0.58 | Medium | Novel prediction |

**Table S11:** Cytoscape result showing protein interacting network of RASSF5

| **Protein Name** | **Degree** | **Average Shortest Path Length** | **Betweenness**  **Centrality** | **Closeness**  **Centrality** |
| --- | --- | --- | --- | --- |
| HRAS (GTPaseHRas) | 20 | 1.33333333 | 0.02887756 | 0.75 |
| KRAS (GTPaseKRas) | 20 | 1.33333333 | 0.02887756 | 0.75 |
| NRAS (GTPaseNRas) | 19 | 1.36666667 | 0.0246782 | 0.73170732 |
| RRAS (Ras-related protein R-Ras) | 18 | 1.4 | 0.03307681 | 0.71428571 |
| RALGDS (Ral guanine nucleotide dissociation stimulator) | 18 | 1.4 | 0.01415094 | 0.71428571 |
| RASSF5 | 18 | 1.4 | 0.13606771 | 0.71428571 |
| RAP1A (Ras-related protein Rap-1A) | 18 | 1.4 | 0.03120347 | 0.71428571 |
| RRAS2 (Ras-related protein R-Ras2) | 17 | 1.43333333 | 0.02880447 | 0.69767442 |
| RAP1B (Ras-related protein Rap-1b) | 17 | 1.43333333 | 0.02739381 | 0.69767442 |
| MRAS (Ras-related protein M-Ras) | 14 | 1.53333333 | 0.02164277 | 0.65217391 |
| RAP1GAP (Rap1 GTPase-activating protein) | 13 | 1.56666667 | 0.01690461 | 0.63829787 |
| MLLT4 (Afadin) | 13 | 1.56666667 | 0.00546438 | 0.63829787 |
| RASSF1 (Ras association domain**-**containing protein1) | 12 | 1.63333333 | 0.04829588 | 0.6122449 |
| STK3 (Serine/Threonine-protein kinase 3) | 7 | 1.86666667 | 0.01054818 | 0.53571429 |
| RAPGEF4 (Rap guanine nucleotide exchange factor 4) | 7 | 1.76666667 | 0.00156178 | 0.56603774 |
| STK4 (Serine/Threonine-protein kinase 4) | 6 | 2.06666667 | 0.00132626 | 0.48387097 |
| SAV1 (Protein Salvador homolog 1) | 6 | 2.06666667 | 0.00132626 | 0.48387097 |
| MOB1A (MOB kinase activator 1A) | 6 | 1.96666667 | 0.0052803 | 0.50847458 |
| MOB1B (MOB kinase activator 1B) | 6 | 1.96666667 | 0.0052803 | 0.50847458 |

**Table S12:** Function of the proteins associated with RASSF5

| **Protein Name** | **Function** |
| --- | --- |
| HRAS (GTPaseHRas) | - Cell growth, differentiation, migration, and apoptosis. |
| KRAS (GTPaseKRas) | - Cell growth, differentiation, migration, and apoptosis. |
| NRAS (GTPaseNRas) | - Cell growth, differentiation, migration, and apoptosis. |
| RRAS (Ras-related protein R-Ras) | - Small GTPase involved in angiogenesis, vascular homeostasis and regeneration. |
| RALGDS (Ral guanine nucleotide dissociation stimulator) | - Effector molecule for Ras protein (R-Ras, H-Ras, K-Ras, and Rap) |
| RASSF5 | - Pro-apoptotic Ras effector. - Promote cell cycle arrest and modulate activity of p53 - Ras regulated tumor suppressor. - Mediate HRAS and KRAS induced apoptosis. |
| RAP1A (Ras-related protein Rap-1A) | - Induces morphological reversion of a cell line transformed by a Ras oncogene. - Counteracts the mitogenic function of Ras. |
| RRAS2 (Ras-related protein R-Ras2) | - GTP-binding protein with GTPase activity. - Might transduce growth inhibitory signals across the cell membrane, exerting its effect shared with Ras proteins but in an antagonistic fashion. |
| RAP1B (Ras-related protein Rap-1b) | - GTP-binding protein that possesses intrinsic GTPase activity. - Plays a role in the establishment of basal endothelial barrier function. |
| MRAS (Ras-related protein M-Ras) | - Cell proliferation. - Weakly activates the MAP kinase pathway. |
| RAP1GAP (Rap1 GTPase-activating protein) | - GTPase activator for the nuclear Ras-related regulatory protein RAP-1A (KREV-1), converting it to the putatively inactive GDP-bound state. |
| MLLT4 (Afadin) | - Role in the organization of epithelial structures of the embryonic ectoderm. - Organization of adherents junctions |
| RASSF1 (Ras association domain**-**containing protein1) | - Potential tumor suppressor. - Required for death receptor-dependent apoptosis. - Activation of STK3/MST2 and STK4/MST1in Fas-induced apoptosis. |
| STK3 (Serine/Threonine-protein kinase 3) | - Stress-activated, pro-apoptotic kinase - Key component of the Hippo signaling pathway - Tumor suppression by restricting proliferation and promoting apoptosis. |
| RAPGEF4 (Rap guanine nucleotide exchange factor 4) | - Guanine nucleotide exchange factor (GEF) for RAP1A, RAP1B and RAP2A small GTPases activated by binding cAMP. |
| STK4 (Serine/Threonine-protein kinase 4) | - Same as STK3 |
| SAV1 (Protein Salvador homolog 1) | - Regulator of STK3/MST2 & STK4/MST1 in the Hippo signaling pathway - Tumor suppression by restricting proliferation and promoting apoptosis. - Promotes cell-cycle exit |
| MOB1A (MOB kinase activator 1A) | - Activator of LATS1/2 in the Hippo signaling pathway - Tumor suppression by restricting proliferation and promoting apoptosis. |
| MOB1B (MOB kinase activator 1B) | - Same as MOB1A |
